# Directional Sensitivity of Cortical Neurons Towards TMS Induced Electric Fields

**DOI:** 10.1101/2023.07.06.547913

**Authors:** Konstantin Weise, Torge Worbs, Benjamin Kalloch, Victor H. Souza, Aurélien Tristan Jaquier, Werner Van Geit, Axel Thielscher, Thomas R. Knösche

## Abstract

We derived computationally efficient average response models of different types of cortical neurons, which are subject to external electric fields from Transcranial Magnetic Stimulation. We used 24 reconstructions of pyramidal cells (PC) from layer 2/3, 245 small, nested, and large basket cells from layer 4, and 30 PC from layer 5 with different morphologies for deriving average models. With these models, it is possible to efficiently estimate the stimulation thresholds depending on the underlying electric field distribution in the brain, without having to implement and compute complex neuron compartment models. The stimulation thresholds were determined by exposing the neurons to TMS-induced electric fields with different angles, intensities, pulse waveforms, and field decays along the somato-dendritic axis. The derived average response models were verified by reference simulations using a high-resolution realistic head model containing several million neurons. Differences of only 1-2% between the average model and the average response of the reference cells were observed, while the computation time was only a fraction of a second compared to several weeks using the cells. Finally, we compared the model behavior to TMS experiments and observed high correspondence to the orientation sensitivity of motor evoked potentials. The derived models were compared to the classical cortical column cosine model and to simplified ball-and-stick neurons. It was shown that both models oversimplify the complex interplay between the electric field and the neurons and do not adequately represent the directional sensitivity of the different cell types.

The derived models are simple to apply and only require the TMS induced electric field in the brain as input variable. The models and code are available to the general public in open-source repositories for integration into TMS studies to estimate the expected stimulation thresholds for an improved dosing and treatment planning in the future.

## 1 Introduction

The extension of current models in the area of transcranial brain stimulation beyond the estimation of the electric fields is elementary to improve our understanding of the underlying stimulation processes. The key question is how the electric field modulates the behavior of neuronal structures. Earlier experimental studies showed that the depolarization threshold of isolated straight axons is inversely proportional to the cosine of the angle between the external current and the nerve fiber (Rushton, 1927). This led to the well known cortical column cosine hypothesis (Fox et al., 2004), assuming that excitable neuronal elements, in particular axons, have a preferential orientation perpendicular to the cortical surface. At first glance, this model seems to be supported by the findings of Rudin and Eisenman (1954) and Ranck (1975), who consistently found that orthodromic currents are more effective than antidromic currents and especially transverse currents. Note, however, that the complex morphology of the neurons does not allow the generalization of observations made in single isolated axons to neuronal populations, because the orientations of the axon segments relative to the external electric field vary and have to be considered statistically. Accounting for these effects requires a model description across multiple scales. This involves first determining the electric field in the brain by solving Maxwell’s equations and then coupling it with detailed mesoscopic neuron models. Aberra et al. (2022) introduced a novel approach to simulate the effects of TMS in head models with morphologically realistic cortical neurons. These authors developed a multi-scale computational model that is capable of quantifying effects of different TMS parameters on the direct response of individual cortical neurons. They created digital representations of neurons that match the geometry and biophysical properties of mature human neocortical cells based on neuronal models of rodent cells from the Blue Brain Project (Markram et al. 2015). These models included a spatial representation of the neuronal compartments as well as experimentally validated electrophysiological parameters (Aberra et al., 2018). They were placed inside the gray matter of a realistic head model and the stimulation thresholds for the generation of action potentials were determined by coupling them with the TMS induced electric fields. The results provide important mechanistic insights into TMS. However, a major limitation of this modeling approach is its high computational cost, which prevents most routine applications of the method in TMS studies. Moreover, a further challenge is that for estimating the overall threshold of a cortical group of neurons, the results of a large number of single simulated responses need to be determined and averaged. This calls for the development of simpler models of neural populations that still accurately account for the modulation of neuronal states through TMS-induced electric fields.

We developed a parsimonious model, which reproduces the effect of the electric field on cortical neurons with high accuracy for different pulse waveforms and geometric electric field parameters. We adapted and extended the approach of Aberra et al. (2020) to derive an average threshold model of layer 2/3 pyramidal cells (L2/3 PC), small, nested, and large basket cells in layer 4 (L4 SBC, L4 NBC, L4 LBC), and layer 5 pyramidal cells (L5 PC). We adapted the pipeline of Aberra et al (2020) in Python and implemented additional improvements and extensions, such as support for SimNIBS 4 and the CHARM head modeling pipeline (Puonti et al. 2020). The code and associated example scripts are published in the open-source Python package *TMS-Neuro-Sim* (https://github.com/TorgeW/TMS-Neuro-Sim). Additionally, we determined estimators for the neuronal recruitment rate, which quantifies the relative number of neurons stimulated by TMS at a given stimulation intensity and field orientation.

To further investigate the derived models, we performed a sensitivity analysis and identified the most influential parameters of the models by determining so-called Sobol indices using a generalized polynomial chaos expansion (Weise et al., 2020b). Moreover, the model was verified by comparing it to results of computationally expensive reference simulations, using a high resolution realistic head model with a large number of realistically shaped neurons located within the motor cortex. Finally, we validated the model by comparing it with TMS experiments by Souza et al. (2022), who intensively investigated the directional sensitivity of motor evoked potentials using a novel multi-coil TMS transducer.

We also compared the results with those of the cortical column cosine (Fox et al., 2004) as well as a simplified ball-and-stick model (Bédard and Destexhe, 2008) adapted for TMS. It turned out that the stimulation properties differ significantly from detailed neurons and that a simplified modeling strategy is not appropriate in this context.

All data and code underlying the results presented in this paper, together with additional details including the average threshold models, the recruitment rate operators, and the neuron compartment models, are publically available in a repository (Weise et al., 2023b), where we provide look-up tables, interpolators, and polynomial approximations for further use.

## 2 Methods

### 2.1 Neuron models

To derive the average neuron response models, we extended the set of neural compartment models by Aberra et al. (2020) from originally five neurons to 24 L2/3 PC, 70 L4 SBC, 70 L4 NBC, 105 L4 LBC, and 30 L5 thick-tufted pyramidal cells (TTPC’s), taken from the Blue Brain Project (Ramaswamy et al., 2015). The cells originate from the somatosensory cortices of P14 male Wistar (Han) rats (Markram et al., 2015). They were stained with biocytin, visually recorded with a bright-field light microscope, and processed by the software Neurolucida (Williston, VT, USA). Shrinkage due to staining in the *z*-axis was corrected during the reconstruction. In an unraveling step, shrinkage in the *xy*-axis was corrected for with a method based on the centered moving window algorithm by smoothing and extending the reach of the branches while maintaining their overall length (Anwar et al., 2009). For branch repair, the cutting planes were first determined and the cut branches were then statistically regrown based on the intact branches. Because some resulting cell morphologies contain impoverished axonal/dendrite branching, a mix-and-match procedure was used to create cells with valid dendrite and axonal reconstructions. As a last step to increase morphological diversity, a cloning procedure was applied. The procedure assigns distributions to branch length and rotation while preserving the overall branching structure.

Because the cells provided were from rats, further modifications were necessary to obtain human-like neurons. We followed the procedure and parameters given in Aberra et al. (2018) to extend the set of neurons. First, the basal dendritic diameter, basal dendritic length, apical dendritic diameter, somatic diameter, and axonal diameter were scaled to create adult human-like neuron morphologies. Second, the axons were myelinated by registering nodes of Ranvier with a width of 1 *μ*m, creating myelinated sections with a length (L) to diameter (D) ratio of L/D=100 and myelinated axon terminals with L/D=70 (Hursh 1939, Hess and Young 1949, Waxman and Kocsis 1995). And third, the ion channel properties were adapted according to the myelination (see Table 1 in Aberra et al., 2018). Fig. 1 provides an overview of the cells used in the study. The average numbers of nodes per cell are 3,541 for L2/3 PC, 14,779 for L4 SBC, 13,091 for L4 NBC, 9,147 for L4 LBC, and 12,514 for L5 PC. For the compartment models, the neurons were discretized with a maximum compartment length of 20 μm. This resulted in an average number of compartments of 766 for L2/3 PC, 1,447 for L4 SBC, 1,762 for L4 NBC, 1,876 for L4 LBC, and 2,008 for L5 PC.

**Figure 1:**
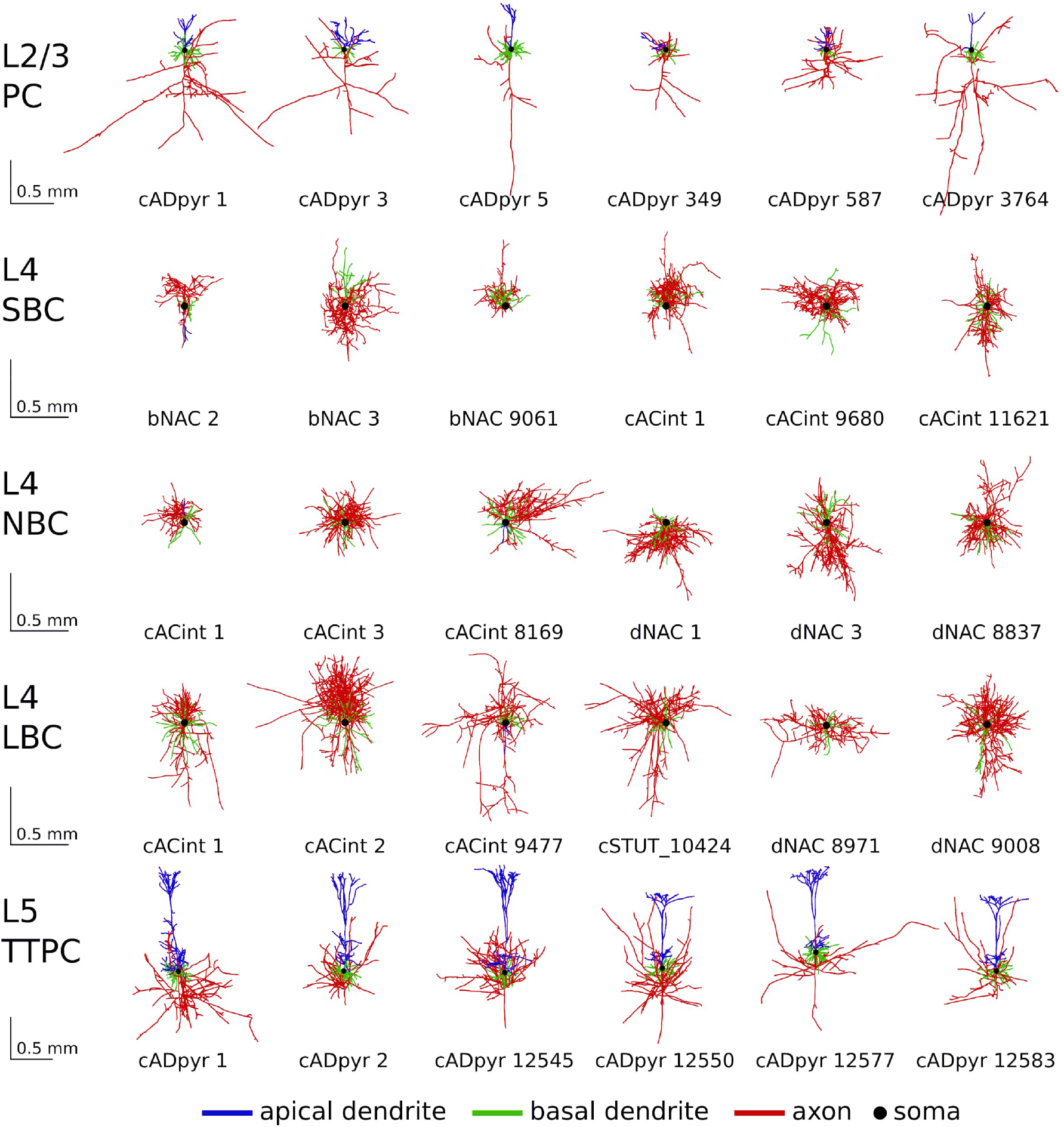
Example morphologies of L2/3 PC, L4 S/N/LBC, and L5 PC: The numbers below the cells indicate the corresponding IDs in the repository Weise et al. (2023b). L4 BC are categorized in small basket cells (SBC), nested basket cells (NBC), and large basket cells (LBC). In total, the study includes 24 L2/3 PC, 70 L4 SBC, 70 L4 NBC, 105 L4 LBC, and 30 L5 thick-tufted pyramidal cells (TTPC’s), taken from the Blue Brain Project (Ramaswamy et al., 2015).

**Table 1:** Sobol indices of the electric field threshold models for L2/3, L4 S/N/LBC, and L5 PC for monophasic and biphasic pulse waveforms.

| Cell type | L2/3 PC |  | L4 SBC |  | L4 NBC |  | L4 LBC |  | L5 PC |  |
| --- | --- | --- | --- | --- | --- | --- | --- | --- | --- | --- |
| wave-form | mono-phasic | bi-phasic | mono-phasic | bi-phasic | mono-phasic | bi-phasic | mono-phasic | bi-phasic | mono-phasic | bi-phasic |
| $\vartheta$ | 0.919 | 0.710 | 0.991 | 0.994 | 0.974 | 0.985 | 0.962 | 0.974 | 0.951 | 0.971 |
| $\Delta\vert \vec{E}\vert $ | 0.052 | 0.247 | 0.001 | 0.001 | 0.001 | 0.001 | 0.003 | 0.002 | 0.037 | 0.017 |
| $\vartheta$ & $\Delta\vert \vec{E}\vert $ | 0.029 | 0.043 | 0.008 | 0.005 | 0.025 | 0.014 | 0.035 | 0.024 | 0.012 | 0.012 |

### 2.2 Coupling of electric fields into neuron models

The electric field **E**(*z, t*) caused by TMS generates an additional extracellular pseudo-potential *φ_e_*(*z, t*). It is coupled into the neurons’ cable equations by integrating the electric field component along the local longitudinal direction **dz** of each neuronal compartment, ranging from, for example, the initial point of a compartment *z_0_* to the end of that compartment *z*:

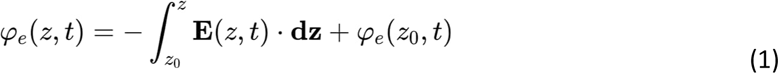

For the realistic head model simulations, the electric field is interpolated to the neurons’ segments using the superconvergent patch recovery approach (Zienkiewicz and Zhu, 1992).

### 2.3 Neuronal simulations

The stimulation behavior of the neurons is analyzed by calculating the transmembrane potential in each compartment using NEURON (Carnevale and Hines, 2006) following a similar approach as Aberra et al. (2020). The spatio-temporal dynamics of the transmembrane potential were modeled according to the Hodgkin-Huxley formalism. Detailed information about the ion channel parameters and membrane time constants can be found in the repository by Weise et al. (2023b) and ModelDB (https://senselab.med.yale.edu/modeldb/ShowModel.cshtml?model=241165) by Aberra et al. (2018). The NEURON simulation was set up with a temperature of 37° C and an initial voltage of -70 mV for each compartment. The simulations were carried out over the course of 1 ms with time steps of 5 µs. The extracellular quasipotentials were scaled by the waveform and the amplitude of the TMS pulse. The used monophasic and biphasic waveforms were taken from a MagPro X100 stimulator (MagVenture A/S, Denmark) with a MagVenture MCF-B70 figure-of-eight coil (P/N 9016E0564) and were recorded using a search coil with a sampling rate of 5 MHz. The recordings were down-sampled to the simulation time steps and normalized to be applicable for scaling the extracellular potential. The cell thresholds are determined as the minimum electric field intensity needed to elicit action potentials in at least three compartments, using a binary search approach with a precision of 0.05 V/m. The simulation environment is implemented and published in the open-source Python package *TMS-Neuro-Sim* (https://github.com/TorgeW/TMS-Neuro-Sim) making use of the Python API of NEURON. The example dataset (Weise et al., 2023b) contains the neuron models together with example scripts detailing the use of the implemented functions.

### 2.4 Average response model of cortical neurons

We exposed the model neurons to electric fields from different directions and strengths to examine their stimulation behavior in detail. We parameterized the electric field direction using spherical coordinates (Fig. 2a). The polar angle *θ* quantifies the angle between the electric field and the somato-dendritic axis (*z*-axis) and ranges from 0° to 180°. The azimuthal angle *φ* quantifies the electric field direction in the horizontal plane perpendicular to the somato-dendritic axis and ranges between 0° and 360°. The coordinate system is defined such that the soma is lying close to the center and the axon extends into negative *z*-direction. Because of the comparatively large extension of the PC from the uppermost dendrites to the lowermost part of the axon, the decay of the electric field along the *z*-axis is not negligible. In simulations of a realistic head model, we found that the electric field can differ up to ±20% per mm over the somato-dendritic axis. A more detailed analysis of the underlying parameter distributions is given later in Section “Sensitivity analysis”. For this reason, we have added an additional parameter to the model, namely the relative change of the electric field magnitude per unit length Δ|**Ẽ**| measured in %/mm.

**Figure 2:**
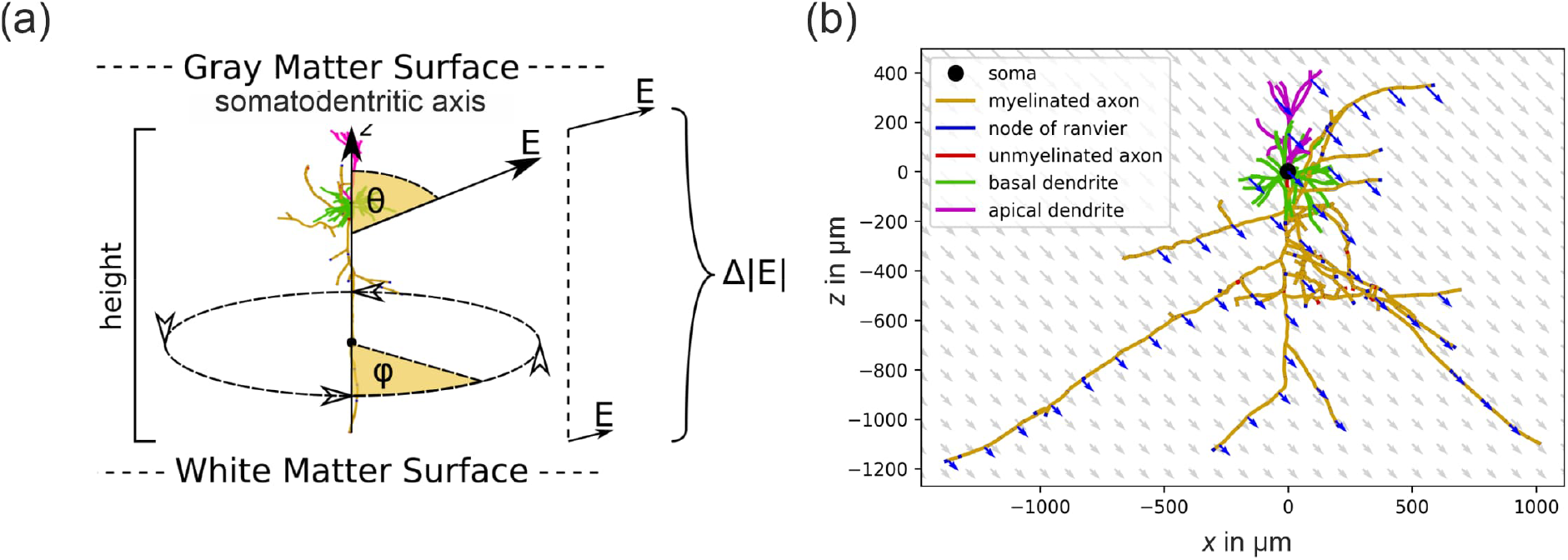
Neurons in an external electric field: (a) Parametrization of the TMS induced electric field relative to cortical neurons; (b) Example of an L2/3 PC, which is exposed to an external electric field with direction *φ*=0°, *θ*=135° and a field decay of Δ|**Ẽ**|=−30%/mm. Note that the electric field is stronger in the upper part of the cell and decreases with depth, as is generally observed in the cortex.

The electric field at each location (*x, y, z*) at firing threshold is then given by:

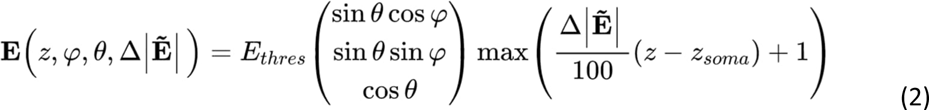

where *z_soma_* is the position of the soma on the *z*-axis (in mm). The soma will have an electric field magnitude of *E_thres_*, which is to be found using the aforementioned binary search approach, while the magnitude for every other section of the cell is linearly interpolated based on their *z*-coordinate. An example of an external electric field distribution with *φ*=0°, *θ*=135° and Δ|**Ẽ**|=−30%/mm is shown in Fig. 2b.

For the derivation of an average response model, all L2/3 and L5 neurons were exposed to an external electric field with a polar angle *θ* (range: [0, 180]°, steps: 3°), an azimuthal angle *φ* (range: [0, 360]°, steps: 6°), and a relative change of the electric field along the somato-dendritic axis Δ|**Ẽ**| (range: [-100, 100] %/mm, steps: 10 %/mm) for both monophasic and biphasic pulse waveforms. After determining the electric field thresholds for each cell for all possible electrical field configurations, an average threshold model was derived by averaging the thresholds over all compartment models and over all azimuthal orientations *φ*, based on the assumption that the spatial locations and tangential orientations of the neurons in the cortex are random.

From the activation thresholds of the individual neurons, we determined the recruitment rate of the neurons in dependence of *θ* and Δ|**Ẽ**|. The recruitment rate estimates the relative number of neurons which were stimulated by TMS at a given stimulation intensity, with zero corresponding to no stimulation and one corresponding to stimulation of all neurons. To this end, we integrated the electric field thresholds along the electric field axis and smoothed the discrete behavior by fitting continuous sigmoid functions of the following type in the least squares sense:

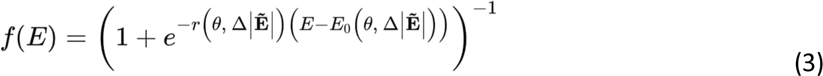

where *E* denotes the electric field, *r*(θ, Δ|**Ẽ**|) the slope, and *E_0_*(θ, Δ|**Ẽ**|) the shift of the sigmoidal functions, which depend on both the polar angle *θ* and the relative change in electric field Δ|**Ẽ**|.

A threshold model was also created for a simplified ball-and-stick neuron model. Typically, ball-and-stick models are used for stimulation by weak electric fields in the context of transcranial electric stimulation (e.g., Aspart et al., 2016) and consist of one segment for the dendrites and one segment for the soma. Because the stimulation thresholds for TMS-induced electric fields of the dendrites are more than 10 times higher than those of the axons, the classical ball-and-stick model had to be modified for TMS. For this purpose, we integrated a straight axon instead of dendrites into the model and determined the stimulation thresholds as a function of the polar angle θ using the approach described by Aberra at al. (2020). For a similar approach, see also the supplementary material of that article). The parameters (axon length and diameter) were adapted such that the thresholds matched those of our complex model.

### 2.5 Sensitivity analysis

A sensitivity analysis of the derived threshold maps was conducted in terms of variations of the electric field parameters *θ* and Δ|**Ẽ**|. We derived a generalized polynomial chaos expansion of the threshold maps using the Python package *pygpc* (Weise et al., 2020b) and determined the first- and second-order Sobol indices that quantify the fraction of the total variance of the threshold that stems from the variance of *θ*, from the variance of Δ|**Ẽ**|, and from a combination of both. The input distributions of both parameters were estimated from the electric field simulation of the high-resolution realistic head model. We extracted the polar angles *θ* and the changes in electric field magnitude Δ|**Ẽ**| in every surface element of layer 5 in the ROI and fitted uniform and beta distributions to the histograms (Fig. 3). We repeated the analysis for layer 2/3 and layer 4, and did not find any major differences in the parameter distributions, due to the close proximity of the layers. For the uncertainty analysis we assumed that both parameters are uncorrelated.

**Figure 3:**
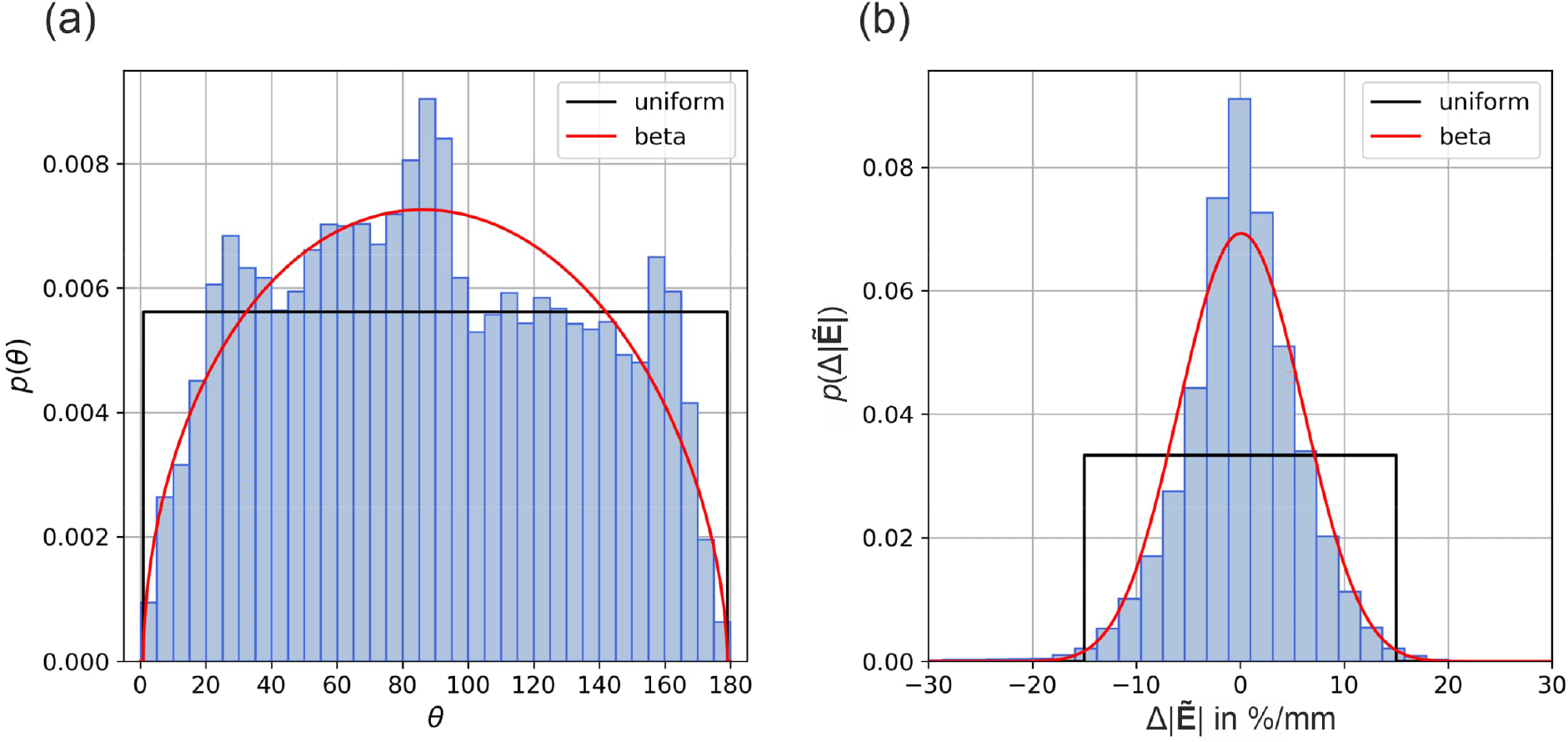
Distribution of electric field parameters on layer 5 in a realistic head model: Histograms and fitted uniform and beta distributions of (a) the polar angle *θ* (uniform parameters: *θ*_min_=0°, *θ*_max_=180°; beta parameters: *θ*_min_=0°, *θ*_max_=180°; *p*=1.51, *q*=1.56) and (b) the relative change of the electric field magnitude Δ|**Ẽ**| (uniform parameters: Δ|**Ẽ**|_min_=-15°, Δ|**Ẽ**|_max_=15°; beta parameters: Δ|**Ẽ**|_min_=-30°, Δ|**Ẽ**|_max_=30°; *p*=13.86, *q*=13.78).

### 2.6 Model verification

In order to verify the average response model, we conducted reference simulations using a high resolution realistic head model, where we explicitly placed the neurons in the ROI and coupled the TMS-induced electric field into them. The head model was created using T1-, T2-, and diffusion weighted MRI. The images were acquired on a 3T MRI scanner (Siemens Skyra) with a 32 channel head coil using the same acquisition parameters as described in Weise et al. (2023a). T1 and T2 images were used for tissue type segmentation. Conductivity tensors in gray and white matter were reconstructed from diffusion weighted images using the volume normalized mapping approach (dwi2cond, https://simnibs.github.io/simnibs/build/html/documentation/command_line/dwi2cond.html, Güllmar et al., 2010). The head model was generated using the headreco pipeline (Nielsen et al., 2018) utilizing SPM12 (https://www.fil.ion.ucl.ac.uk/spm/software/spm12/, Penny et al., 2011) and CAT12 (http://www.neuro.uni-jena.de/cat/, Gaser et al., 2021). A region of interest (ROI) was defined around the handknob area (FreeSurfer, (http://surfer.nmr.mgh.harvard.edu/, Fischl et al., 1998; Dale et al., 1999) based on the fsaverage template. This covered parts of somatosensory cortex (BA1, BA3), primary motor cortex M1 (BA4), and dorsal premotor cortex PMd (BA6). The head model was refined in the ROI to provide accurate electric field values for the neuron models (Fig. 4). The final head model is composed of ∼5·10^6^ nodes and ∼29·10^6^ tetrahedra. The tetrahedra in the ROI have an average edge length of 0.45 mm and an average volume of 0.01 mm³. The model consists of six tissue types with the following electrical conductivities: white matter (0.126 S/m), grey matter (0.275 S/m), cerebrospinal fluid (1.654 S/m), bone (0.01 S/m), skin (0.465 S/m), and eyeballs (0.5 S/m) (Thielscher et al., 2015; Wagner et al., 2004). The entire process from MRI acquisition to the final head model can be reproduced in detail using the protocol of Weise et al. (2023a) (steps 1-20) and details of the FEM are given in Saturnino et al. (2019).

**Figure 4:**
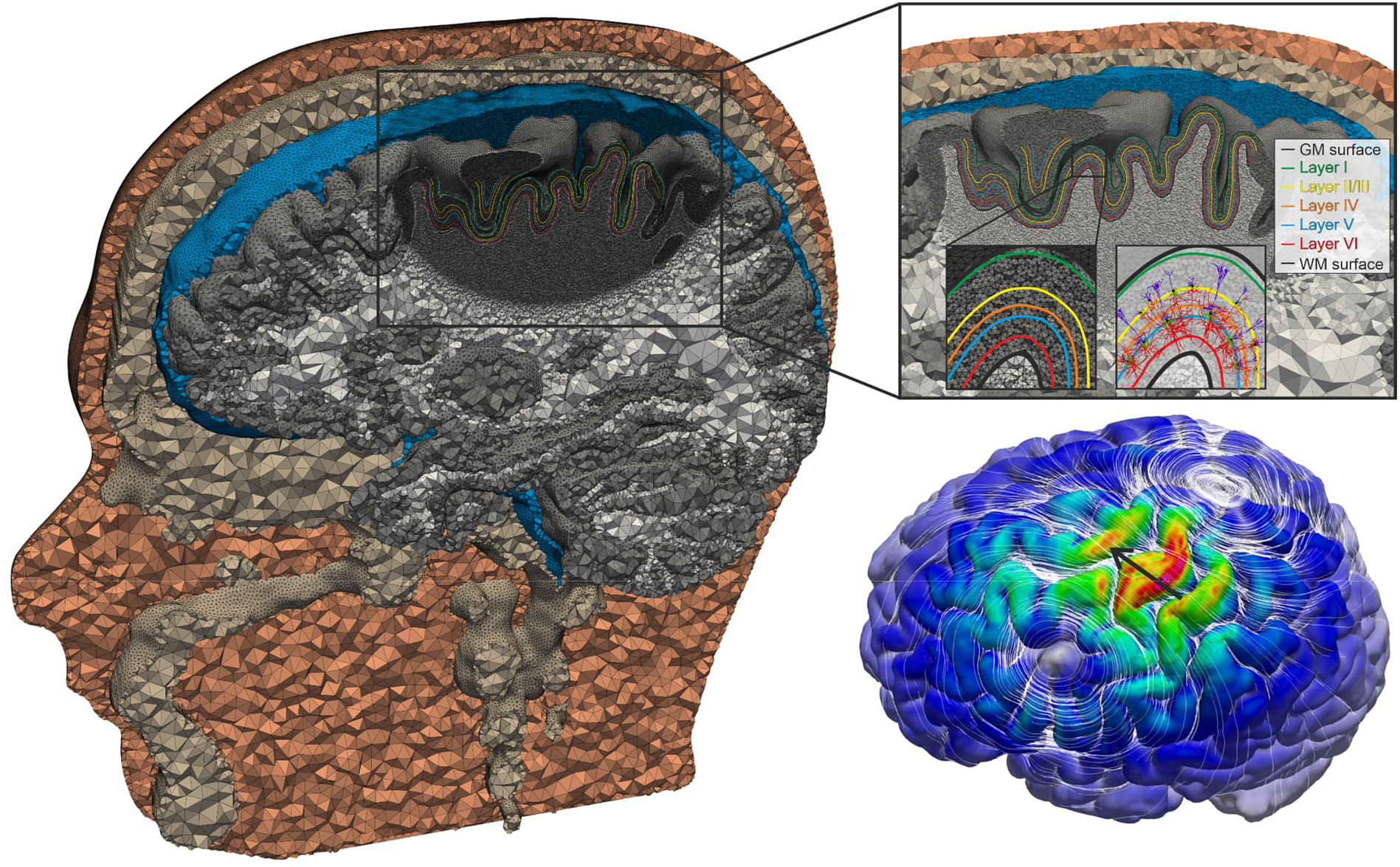
Realistic head model with cortical layers and neurons: The model was constructed with SimNIBS v3.2.6 (Thielscher et al. 2015) using *headreco* (Nielsen et al., 2018). In the M1 region, the cortical layers 1-6 are generated and the mesh is refined to ensure highly accurate electric field profiles, which are coupled into the compartment models of cortical neurons. The bottom right inset shows an example of the magnitude of the electric field as color code and its orientation as white streamlines. The black arrow indicates the coil orientation.

In order to place the neurons at the right locations in the cortex, we added cortical layers to the head model. The normalized depths of the six cortical layers range between 0 (gray matter surface, i.e. pia mater) and 1 (white matter surface) and were estimated from primate motor cortex slices (García-Cabezas et al., 2014). The normalized depths of the layer centers are 0.06 for layer 1, 0.4 for layer 2/3, 0.55 for layer 4, 0.65 for layer 5, and 0.85 for layer 6 (Aberra et al., 2020). We linearly interpolated between the white matter surface (1) and the gray matter surface (0) using the vertex positions of the two surfaces. To extract the cortical layers as isosurfaces from the 3D interpolation, we used a marching cubes algorithm (Fig. 3) (Lorensen et al., 1987). In every ROI surface element (size ∼1 mm²), we placed all cells and rotated them from 0° to 360° in steps of 6°. This resulted in a total number of 12,947,040 L2/3 PC, 130,080,300 L4 S/N/LBC, and 15,760,800 L5 PC in the ROI to simulate. The total pure simulation time using 48 cluster nodes with 72 cores each (Intel Xeon Platinum 8360Y, 256 GB RAM) was approximately 40 days for both monophasic and biphasic waveforms.

The electric field calculations were conducted using SimNIBS v3.2.6 (Thielscher et al, 2015; Saturnino et al., 2019) using a regular figure-of-eight coil (MCF-B65, Magventure, Farum, Denmark), which is placed over the M1 region with an orientation of 45° towards the *fissura longitudinalis*. The angle *θ* of the electric field was calculated with respect to the surface normal of the cortical layers in the ROI. Likewise, the percentage change of the electric field magnitude between the WM and GM surfaces Δ|**Ẽ**| was calculated by extracting the field at a normalized depth of 10 % of the distance between the current layer and the WM and GM surface, respectively, in order to avoid numerical inaccuracies close to the tissue boundaries. The simulation time of the electric field was relatively short compared to the NEURON simulations and took a few seconds.

### 2.7 Experimental validation

We compared the derived recruitment models to experimental observations from Souza et al. (2022). TMS experiments were conducted to investigate the orientation selectivity of neuronal excitability using a novel two-coil multi-channel TMS transducer for manipulating the electric field orientation. The advantage of the used two-coil TMS transducer is the possibility to precisely manipulate the pulse orientation electronically with high accuracy (∼1°), without physically moving the transducer. To measure the effect of the electric field orientation on the motor evoked potential (MEP) amplitude, five single TMS pulses were applied to the *abductor pollicis brevis* (APB) muscle hotspot at each of 120 different pulse orientations (0–360°; steps of 3°) with a stimulation intensity of 110% of the resting motor threshold (rMT). The MEPs from the APB muscle were recorded from 11 subjects (mean age: 30 years, range 24-41; four women) using surface electromyography electrodes with a belly-tendon montage. TMS pulses had a trapezoidal monophasic waveform (timings: 60 µs of rising, 30 µs of hold, 43.2 µs of falling) and were delivered using a custom power electronics. The interstimulus intervals were pseudo-randomized following a uniform distribution between 4 and 6 s. In two other subjects (ages 31 and 36 years; two men; right-handed), the experiment was repeated with stimulation intensities of 110%, 120%, 140%, and 160% rMT. The order of the orientations and intensities of the pulses was pseudo-randomized. A detailed description of the experimental procedure and TMS hardware is given in Souza et al. (2022).

## 3 Results

### 3.1 Stimulation behavior of L2/3 PCs

The results of the average response model of L2/3 PCs in case of a monophasic TMS pulse is shown in Fig. 5. In Fig. 5a, the electric field thresholds are shown as function of the polar angle *θ* and the relative change in electric field magnitude Δ|**Ẽ**|. For parameter combinations of particular interest, we illustrated the stimulation location on a representative neuron. Lowest thresholds can be observed when the electric field is parallel to the somato-dendritic axis. This effect is enhanced for positive electric field changes, that is, when the electric field increases from the dendrites to the lower parts of the axons.

**Figure 5:**
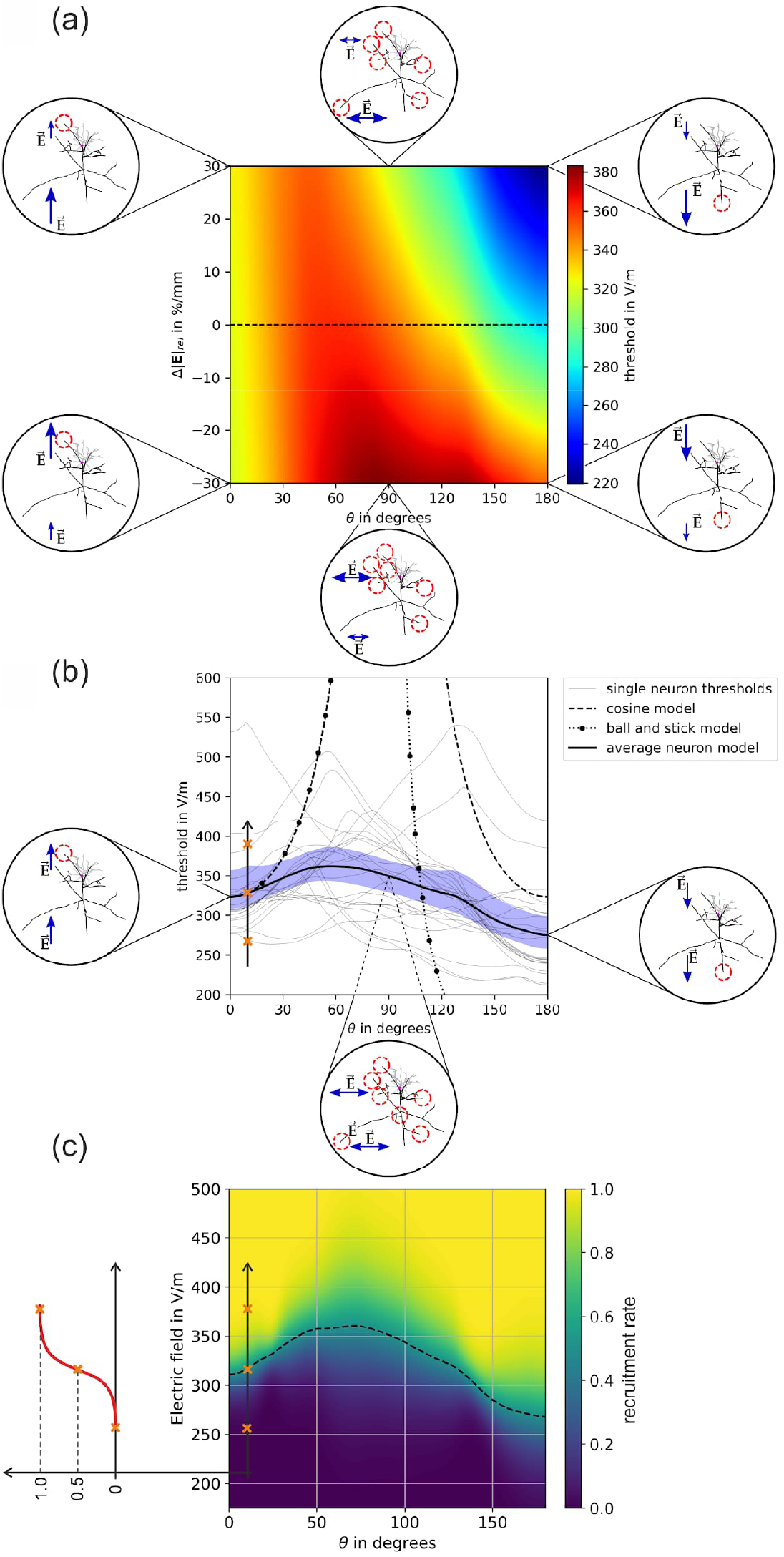
Stimulation behavior of L2/3 PCs for monophasic excitation: (a) Threshold map in dependence of the polar angle *θ* and the relative change of the electric field over the somato-dendritic axis Δ|**Ẽ**|. The insets show the locations of excitation, the red circles indicate the activated terminals. Blue arrows indicate the electric field direction and magnitude; (b) Thresholds of individual neurons for Δ|**Ẽ**|=0 %/mm along the dashed line in (a). The blue area shows the 95th percentile of the confidence interval of the mean. The equivalent cortical column cosine model is *y*(θ) = ŷ|cos (θ)|^-1^ with ŷ=323.27 V/m (dashed line);; the axon parameters of the equivalent ball-and-stick model are *l* = 660 μm and *d*= 15μm (dotted line); (c) Recruitment rate for Δ|**Ẽ**|=0 %/mm derived from the individual neuron activation in (b) by integrating over the electric field thresholds. The dashed line indicates the electric field intensity where the recruitment rate is 0.5.

The behavior of the 24 individual L2/3 neurons is shown in Fig. 5b for homogeneous electric fields, i.e., Δ|**Ẽ**|=0 %/mm (dashed line in Fig. 5a). It can be observed that the electric field thresholds are highest when the electric field is approximately normal to the somato-dendritic axis (*θ*≈90°). Since in this case the electric field can approach from all azimuthal directions *φ* over which the average was taken, there are several potential stimulation sites. The thresholds are decreasing again when the electric field rotates further until it is pointing antidromic, i.e. from the axons to the dendrites (*θ*=0°). In this case the activation takes place at cortico-cortical axon branches pointing upwards. This effect is stable in terms of electric field changes along the somato-dendritic axis. It is noted that due to the geometrical relations of the two electric field angles *θ* and *φ*, the more parallel the electric field is to the somato-dendritic axis (*θ*→0°, *θ*→180°) the less the azimuthal direction *φ* plays a role on the stimulation behavior of the neurons. The thresholds for tangential electric fields (*θ*=90°) are about 17% higher compared to normal electric fields (*θ*=0° and *θ*=180°). The recruitment rates are shown in Fig. 5c for homogeneous electric fields, i.e. Δ|**Ẽ**|=0 %/mm.

We provide the data of the threshold map from Fig. 5a, the individual neuron behavior from Fig. 5b and beyond, as well as the data of the recruitment rate from Fig. 5c in the associated dataset (Weise et al. 2023b). In addition, we provide Python based SciPy interpolators (Virtanen et al., 2020), whose simple usage is also explained in attached scripts.

The results for biphasic excitation are shown in Fig S1 in the *Supplemental Material*.

### 3.2 Stimulation behavior of L4 BCs

#### 3.2.1 Small Basket cells

The results of the average response model of L4 SBCs in case of a monophasic excitation is shown in Fig. 6. A pronounced directional sensitivity can be observed also for this cell type. Again, lowest thresholds can be observed when the electric field is parallel to the somato-dendritic axis (*θ*=0° and *θ*=180°). The thresholds are about 17% higher when the external electric field is tangential to the cells (*θ*=90°). The thresholds are slightly affected if the electric field changes along the somato-dendritic axis (Δ|**Ẽ**|≶0 %/mm). Compared to other cells, the average threshold is about 20% and 46% higher for L4 SBCs than for L2/3 PCs and L5 PCs, respectively. The results for biphasic excitation are shown in Fig S2 in the *Supplemental Material*.

**Figure 6:**
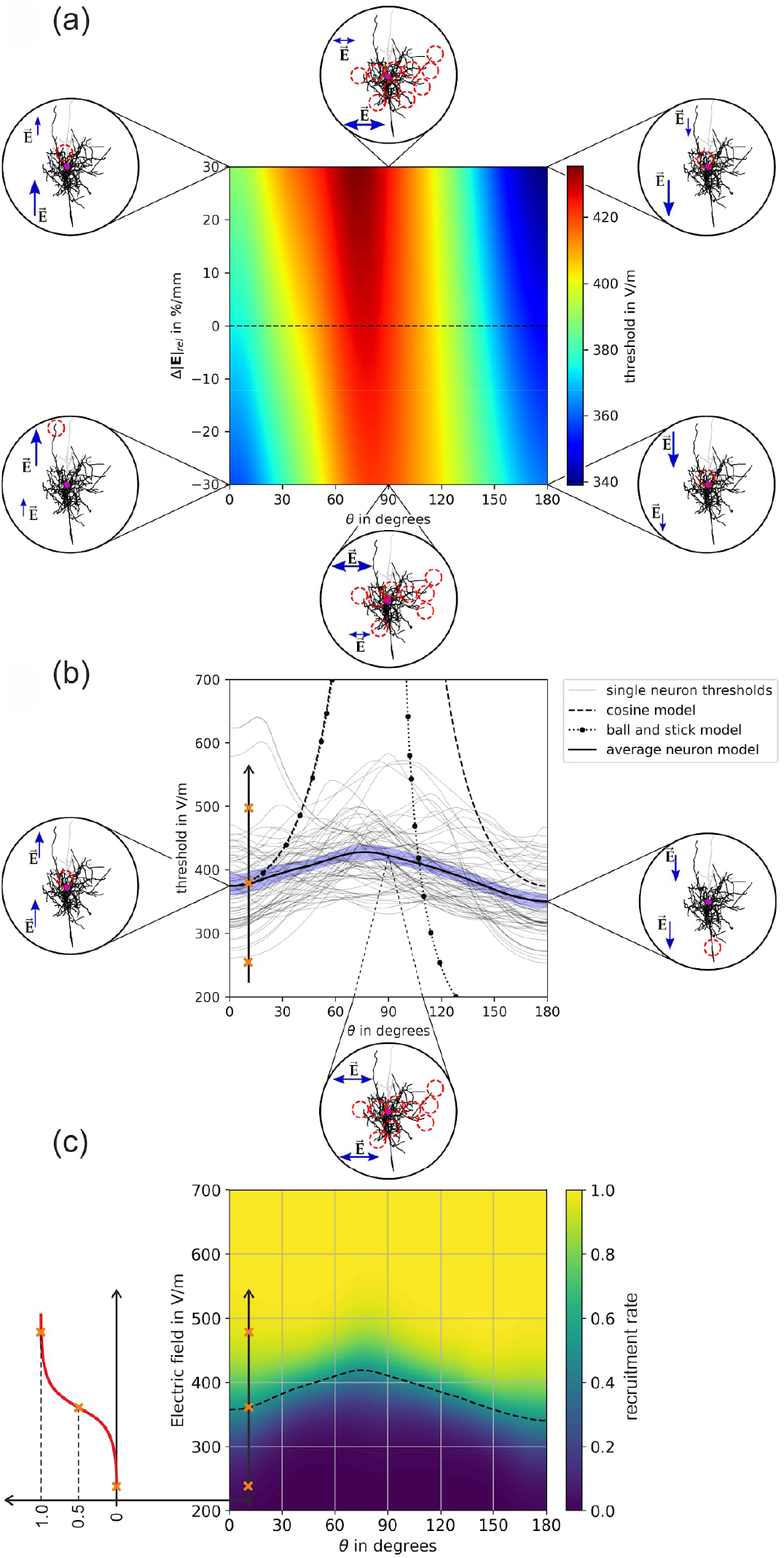
Stimulation behavior of L4 SBCs for monophasic excitation: (a) Threshold map in dependence of the polar angle *θ* and the relative change of the electric field over the somato-dendritic axis Δ|**Ẽ**|. The insets show the locations of excitation, the red circles indicate the activated terminals. Blue arrows indicate the electric field direction and magnitude; (b) Thresholds of individual neurons for Δ|**Ẽ**|=0 %/mm along the dashed line in (a). The blue area shows the 95th percentile of the confidence interval of the mean. The equivalent cortical column cosine model is *y*(*θ*) = *ŷ*|cos(*θ*)|^−1^ with ŷ=178.43 V/m (dashed line); the axon parameters of the equivalent ball-and-stick model are and (dotted line); (c) Recruitment rate for Δ|**Ẽ**|=0 %/mm derived from the individual neuron activation in (b) by integrating over the electric field thresholds. The dashed line indicates the electric field intensity where the recruitment rate is 0.5.

#### 3.2.2 Nested Basket cells

The results of the average response model of L4 NBCs in case of a monophasic excitation is shown in Fig. 7. Their axonal arborization is distinct from pyramidal cells because they form intricate networks of branches that wrap around the soma of nearby pyramidal cells, forming a characteristic “basket” structure. Their axonal structure is generally more isotropic compared to pyramidal cells or SBCs and LBCs. Nevertheless, the thresholds for tangential electric fields are about 14% higher compared to normal electric fields (*θ*=0° and *θ*=180°). On average, the thresholds of L4 NBCs are 2% and 23% higher compared to L2/3 PCs and L5 PCs, respectively. The results for biphasic excitation are shown in Fig S3 in the *Supplemental Material*.

**Figure 7:**
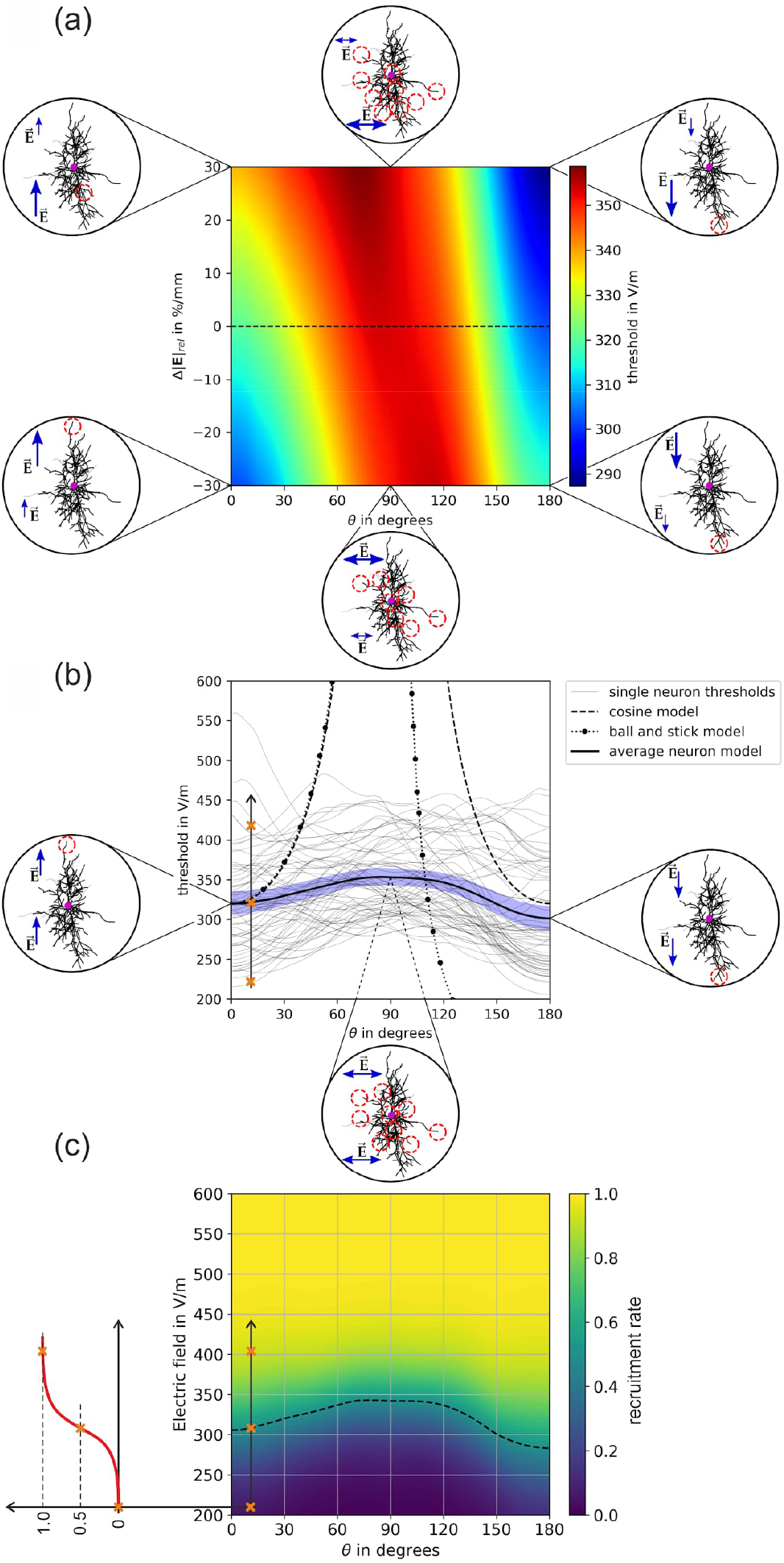
Stimulation behavior of L4 NBCs for monophasic excitation: (a) Threshold map in dependence of the polar angle *θ* and the relative change of the electric field over the somato-dendritic axis Δ|**Ẽ**|. The insets show the locations of excitation, the red circles indicate the activated terminals. Blue arrows indicate the electric field direction and magnitude; (b) Thresholds of individual neurons for Δ|**Ẽ**|=0 %/mm along the dashed line in (a). The blue area shows the 95th percentile of the confidence interval of the mean. The equivalent cortical column cosine model is *y*(*θ*) = *ŷ*|cos(*θ*)|^−1^ with ŷ=178.43 V/m (dashed line); the axon parameters of the equivalent ball-and-stick model are and (dotted line); (c) Recruitment rate for Δ|**Ẽ**|=0 %/mm derived from the individual neuron activation in (b) by integrating over the electric field thresholds. The dashed line indicates the electric field intensity where the recruitment rate is 0.5.

#### 3.2.3 Large Basket cells

The threshold results of L4 LBCs for monophasic excitation are shown in Fig. 8. Compared to PCs, LBCs exhibit a high degree of collateralization in their axonal tree. They can have multiple branches and collaterals that extend in different directions within the same cortical layer or across layers. A distinct directional sensitivity of the thresholds can be again observed together with an asymmetric modulation when the electric field changes along the somato-dendritic axis. On average, the thresholds of L4 LBCs are 9% lower than L2/3 PCs and 11% higher compared to L5 PC, respectively. Of all the basket cells investigated, the LBCs have the lowest thresholds. The average thresholds of LBCs are 24% and 10% lower compared to SBCs and NBCs, respectively. The results for biphasic excitation are shown in Fig S4 in the *Supplemental Material*.

**Figure 8:**
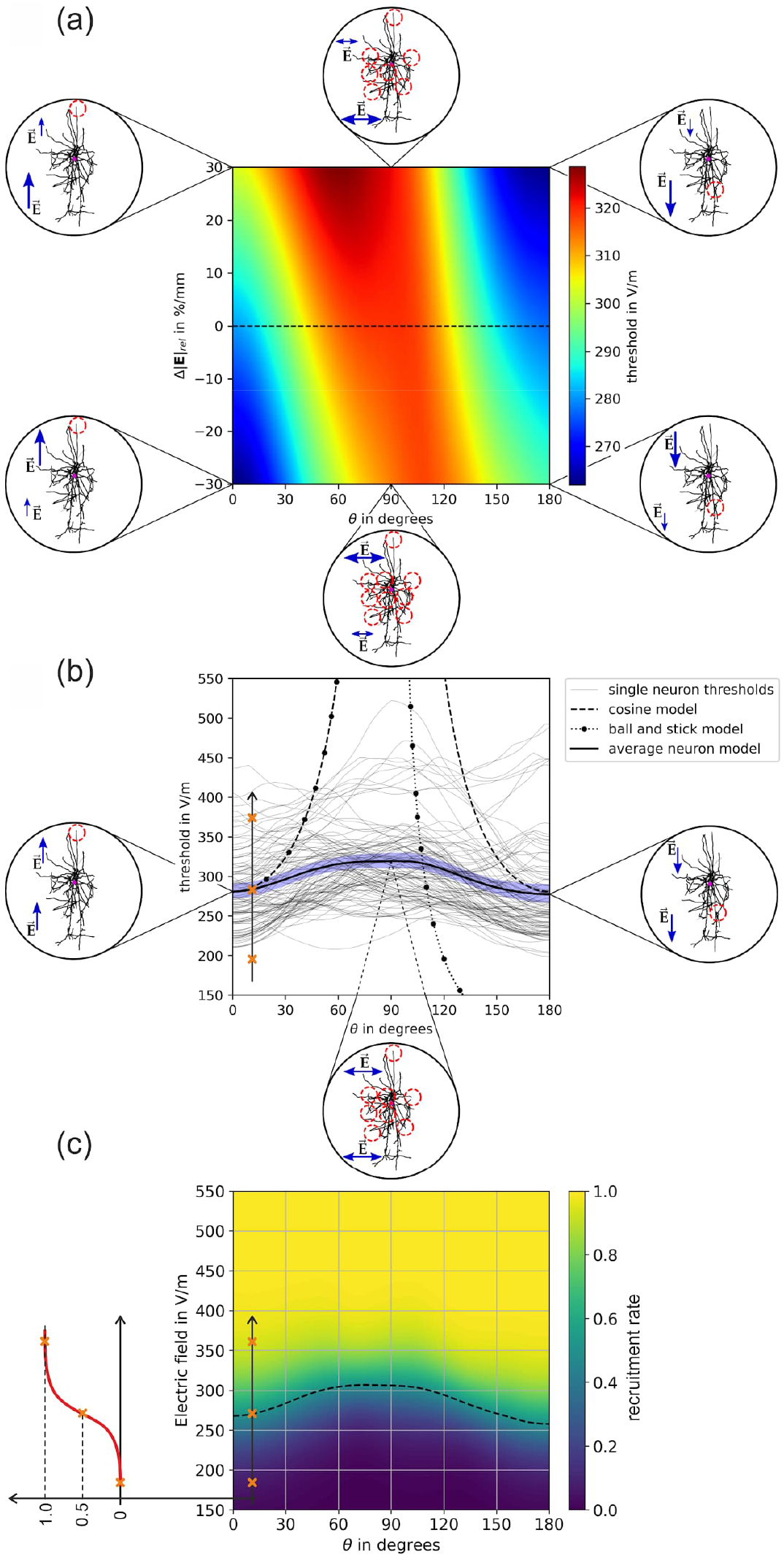
Stimulation behavior of L4 LBCs for monophasic excitation: (a) Threshold map in dependence of the polar angle *θ* and the relative change of the electric field over the somato-dendritic axis Δ|**Ẽ**|. The insets show the locations of excitation, the red circles indicate the activated terminals. Blue arrows indicate the electric field direction and magnitude; (b) Thresholds of individual neurons for Δ|**Ẽ**|=0 %/mm along the dashed line in (a). The blue area shows the 95th percentile of the confidence interval of the mean. The equivalent cortical column cosine model is *y*(*θ*) = *ŷ*|cos(*θ*)|^−1^ with ŷ=178.43 V/m (dashed line); the axon parameters of the equivalent ball-and-stick model are and (dotted line); (c) Recruitment rate for Δ|**Ẽ**|=0 %/mm derived from the individual neuron activation in (b) by integrating over the electric field thresholds. The dashed line indicates the electric field intensity where the recruitment rate is 0.5.

### 3.3 Stimulation behavior of L5 PCs

The stimulation behavior for L5 PCs in case of monophasic excitation is shown in Fig. 9. Coto the behavior of the other cell types investigated, the L5 PCs have the lowest average thresholds (Fig 9a). The thresholds for tangential electric fields (*θ*=90°) are about 15% higher compared to normal electric fields (*θ*=0° and *θ*=180°). The results of the individual neurons in Fig. 9b show that the variance of the thresholds increases with increasing *θ*. At *θ*=180°, a cluster of neurons can be identified that have very low stimulation thresholds. These are paralleled by a few neurons that have very high stimulation thresholds compared to this group. This affects the recruitment rate in Fig. 9c, whose 50% level (dashed line) is lower at *θ*=180° than at *θ*=0°. The most efficient way to stimulate L5 PCs is the application of electric fields with a polar angle of *θ*=0° and a negative change in electric field across the somato-dendritic axis (Δ|**Ẽ**|<0) or by applying electric fields with an angle of *θ*=180° together with a positive field change (Δ|**Ẽ**|>0). For *θ*=180° the stimulation locations are at the lower axons indicating a tendency for cortico-spinal activation. In contrast, when the electric field is antidromic at *θ*=0°, axon collaterals in the upper part are preferentially stimulated, indicating cortico-cortical activation of, for example, connected populations of L2/3 PCs. The stimulation behavior is much more diverse for transverse electric fields around *θ*≈90° due to the variety of azimuthal angles *φ* in which cortico-cortical and cortico-spinal connections can be stimulated. The results for biphasic excitation are shown in Fig S5 in the *Supplemental Material*.

**Figure 9:**
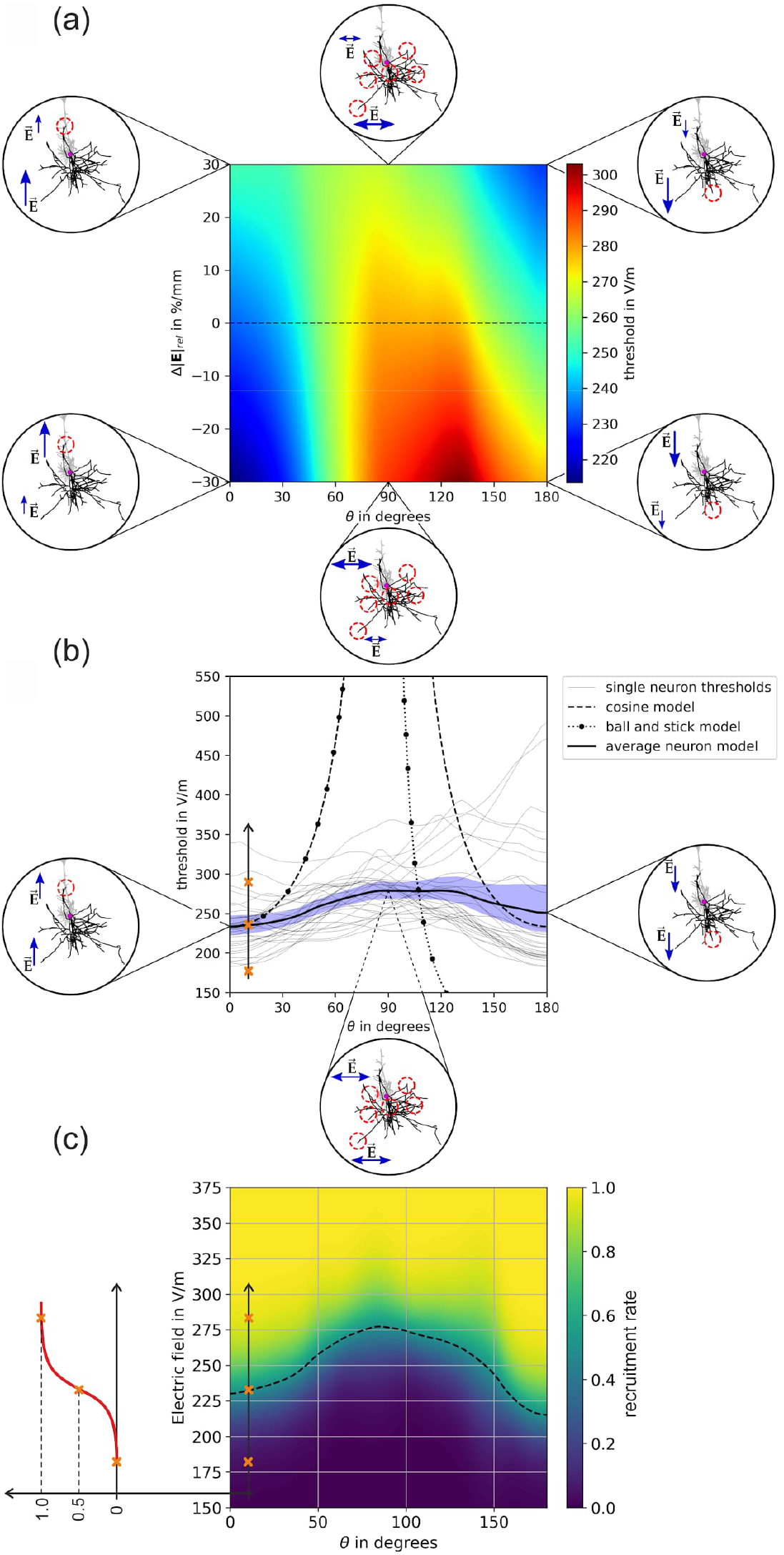
Stimulation behavior of L5 PCs for monophasic excitation: (a) Threshold map in dependence of the polar angle *θ* and the relative change of the electric field over the somato-dendritic axis Δ|**Ẽ**|. The insets show the locations of excitation, the red circles indicate the activated terminals. Blue arrows indicate the electric field direction and magnitude; (b) Thresholds of individual neurons for Δ|**Ẽ**|=0 %/mm along the dashed line in (a). The blue area shows the 95th percentile of the confidence interval of the mean. The equivalent cortical column cosine model is *y*(*θ*) = *ŷ*|cos(*θ*)|^−1^ with ŷ=233.66 V/m (dashed line); the axon parameters of the equivalent ball-and-stick model are and (dotted line); (c) Recruitment rate for Δ|**Ẽ**|=0 %/mm derived from the individual neuron activation in (b) by integrating over the electric field thresholds. The dashed line indicates the electric field intensity where the recruitment rate is 0.5.

### 3.4 Recruitment order and relative threshold ranges

For each cell type investigated, different stimulation thresholds were observed depending on the polar angle *θ* and the relative change of the electric field over the somato-dendritic axis Δ|**Ẽ**|. In Fig. 10, an overview of the threshold ranges of all investigated cell types relative to the mean of L5 PCs, is shown, assuming a constant electric field along the somatodendritic axis (Δ|**Ẽ**|=0 %/mm). It is evident that L5 PCs are recruited first due to their relatively low thresholds. The L4 LBCs have the second lowest thresholds followed by the L2/3 PCs and the L4 NBCs. The small basket cells are directly stimulated only at higher stimulation intensities. An analogous observation was also made for biphasic TMS pulses and the results are reported in Fig. S6 in the *Supplemental Material*.

**Figure 10:**
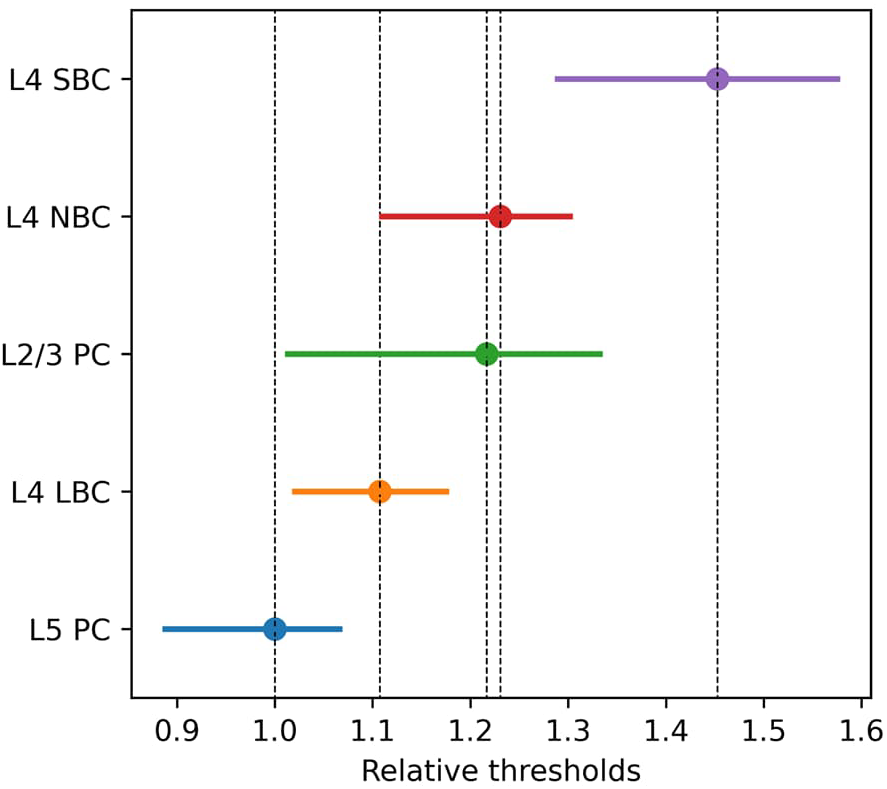
Recruitment order and relative threshold ranges of pyramidal and basket cells for monophasic TMS excitation: Threshold ranges of all investigated cell types relative to the mean of L5 PCs, is shown assuming a constant electric field along the somatodendritic axis (Δ|**Ẽ**|=0 %/mm). The dots indicate the mean thresholds and the ranges stem from the variability across the polar angle *θ* from 0° to 180°.

### 3.5 Sensitivity analysis

We used a 15th order approximation to construct the surrogate models of the threshold maps using *pygpc* (Weise et al., 2020b). The normalized root mean square deviation between the original model and the gPC approximation is 0.32% for L5 PC under monophasic excitation derived from 10^5^ random samples. The accuracies of the gPC approximations of the L2/3 and L4 cells are very similar and given in the repository by Weise et al. (2023b). The results of the sensitivity analysis of the threshold map of L5 PC with monophasic excitation is shown in Fig. 11. It can be seen that the surrogate model (Fig. 11b) almost perfectly resembles the behavior of the original model (Fig. 11a). The absolute differences between both is shown in Fig. 11c. The probability density distribution of the electric field threshold is shown in Fig. 11d under the assumption that the parameters *θ* and Δ|**Ẽ**| are beta distributed as in case of the realistic head model simulations (see Fig. 3 for parameter values). It can be observed that the distribution is u-shaped and bimodal because of the cyclic behavior of the electric field threshold over the polar angle *θ*. The results for L5 in case of monophasic excitation as well as for L2/3 PC, and L4 S/N/LBC for both monophasic and biphasic excitation are given in the repository Weise et al. (2023b).

**Figure 11:**
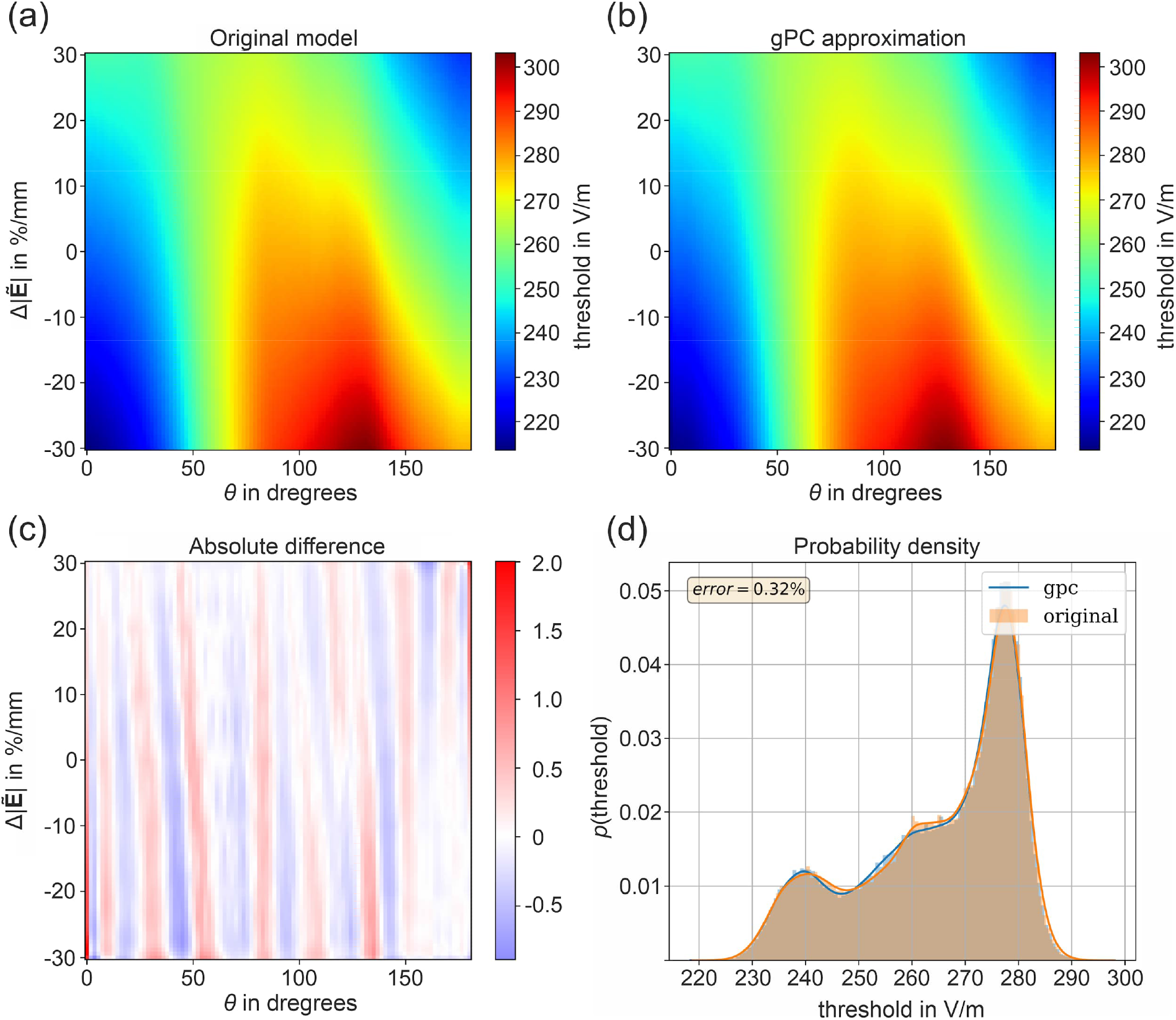
Results of the sensitivity analysis of the electric field threshold map of L5 PCs with monophasic excitation: (a) Original model of the threshold map of L5 PC with monophasic excitation; (b) gPC approximation (surrogate) of the original model; (c) Absolute difference between the original model and the gPC approximation; (d) Probability density of the electric field threshold for the original model and the gPC approximation using N=10^5^ samples under the assumption that *θ* and Δ|**Ẽ**| are beta distributed (see Fig. 3 for parameters).

The Sobol indices, i.e. the fractions of the total variance, which originate from *θ*, Δ|**Ẽ**|, and the combination of both are given in Table 1. The polar angle *θ* has the strongest influence on the stimulation behavior for all investigated cell types. In contrast, the Sobol indices of Δ|**Ẽ**| are much lower, ranging between 2-5%, but the parameter significantly contributes to the increase of the accuracy of the overall model. There is even an exception in the L2/3 cells under biphasic excitation, where the influence reaches almost 25%.

### 3.6 Verification

The average threshold model is compared against reference simulations using a high resolution realistic head model. For the application of the average model, we extracted the electric field parameters *θ* and Δ|**Ẽ**| in every cortical element in the ROI on layer 2/3, 4, and 5 and determined the electric field thresholds (Fig. 5-9) by linearly interpolating the data between the sampling points. The approach is computationally very efficient, as the computation time is only a fraction of the one needed for the electric field computation. In the reference simulations, we calculated the stimulation thresholds for every neuron at every cortical location in the ROI separately by coupling the actual electric fields from the realistic head model into every neural compartment. Finally we averaged the thresholds over all neurons and assigned the resulting average threshold to the ROI element. The resulting electric field threshold maps between the average threshold model and the reference simulations are shown in Fig. 12 for all cell types under investigation. The results for biphasic excitation are shown in Figure S8 in the *Supplemental Material*.

**Figure 12:**
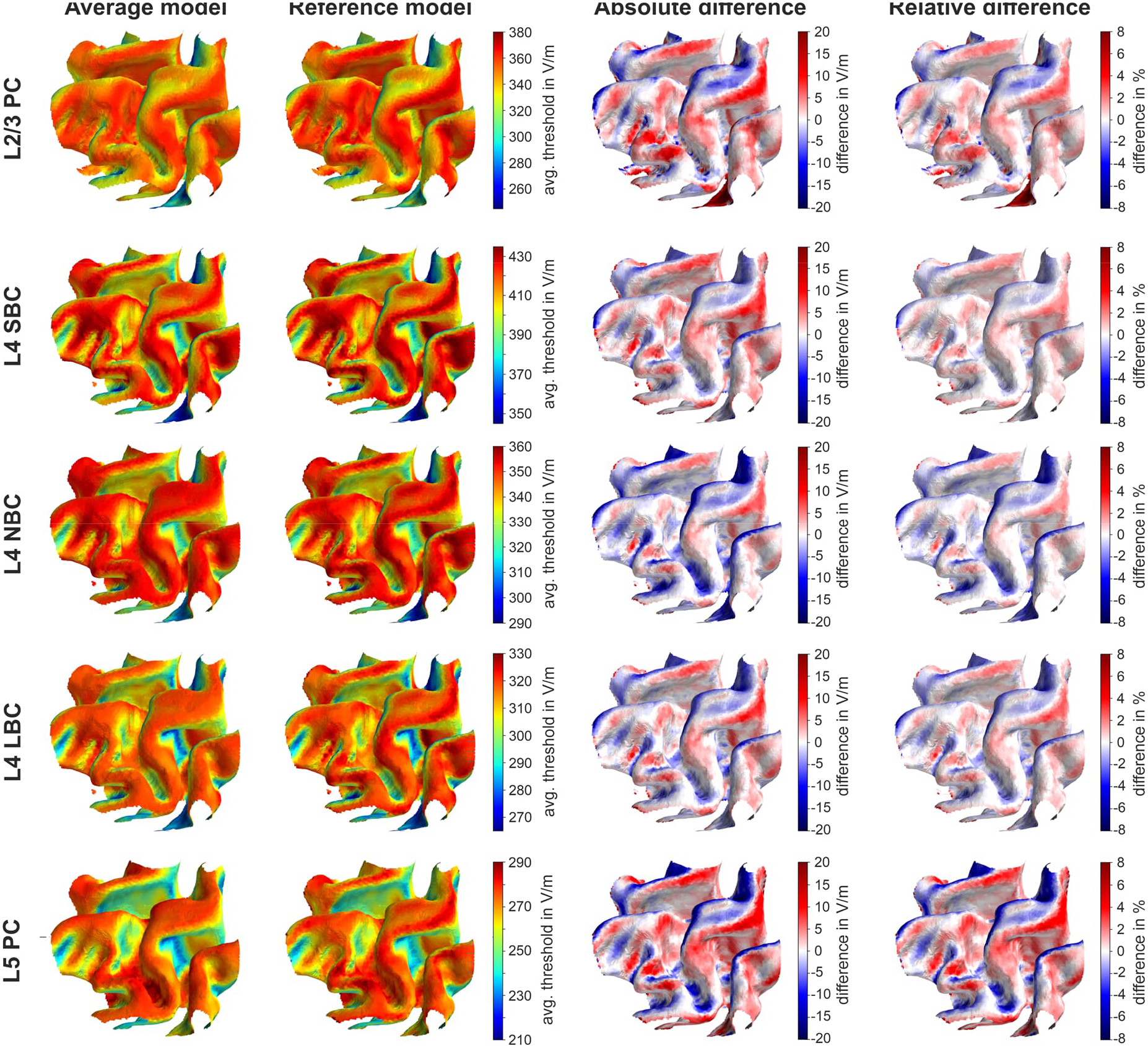
Comparison of electric field threshold maps (in V/m) for monophasic excitation determined using the average model and the reference model: The rows show the electric field threshold maps (in V/m) of the L2/3 PC, L4 S/N/LBC and the L5 PC between the average model (first column) and the reference model (second column). The last two columns show the absolute and relative difference between the models. The underlying electric field distribution and field direction is shown in Fig. 4. The results for biphasic excitation are shown in Figure S8.

For all stimulation conditions, the two models agree very well. The highest relative differences are in the range of ±8% and can be observed mainly at the gyral rims and the sulcal walls. Comparing the distributions and signs of the relative differences between monophasic and biphasic waveforms, it can be observed that they slightly depend on the stimulation waveform and the resulting current direction.

Additionally, we calculated the stimulation threshold maps when the TMS coil is located over the M1 region with a 45° orientation towards the *fissura longitudinalis*. For this, we determined the ratio between the electric field threshold map from Fig. 12 and the corresponding electric field distribution of this particular coil position, which was calculated assuming a normalized stimulation strength of 1 A/µs. This results in a map of the stimulation strength of the TMS stimulator in A/µs needed to stimulate the neurons with this particular coil position. Again a high agreement between the average threshold model and the reference model can be observed. Note that the relative difference distributions in the last column of Fig. 13 are the same as for the electric field threshold maps from Fig. 12 since the electric field distribution is cut out in the error calculation. The analogous results for biphasic excitation are shown in Fig. S9.

**Figure 13:**
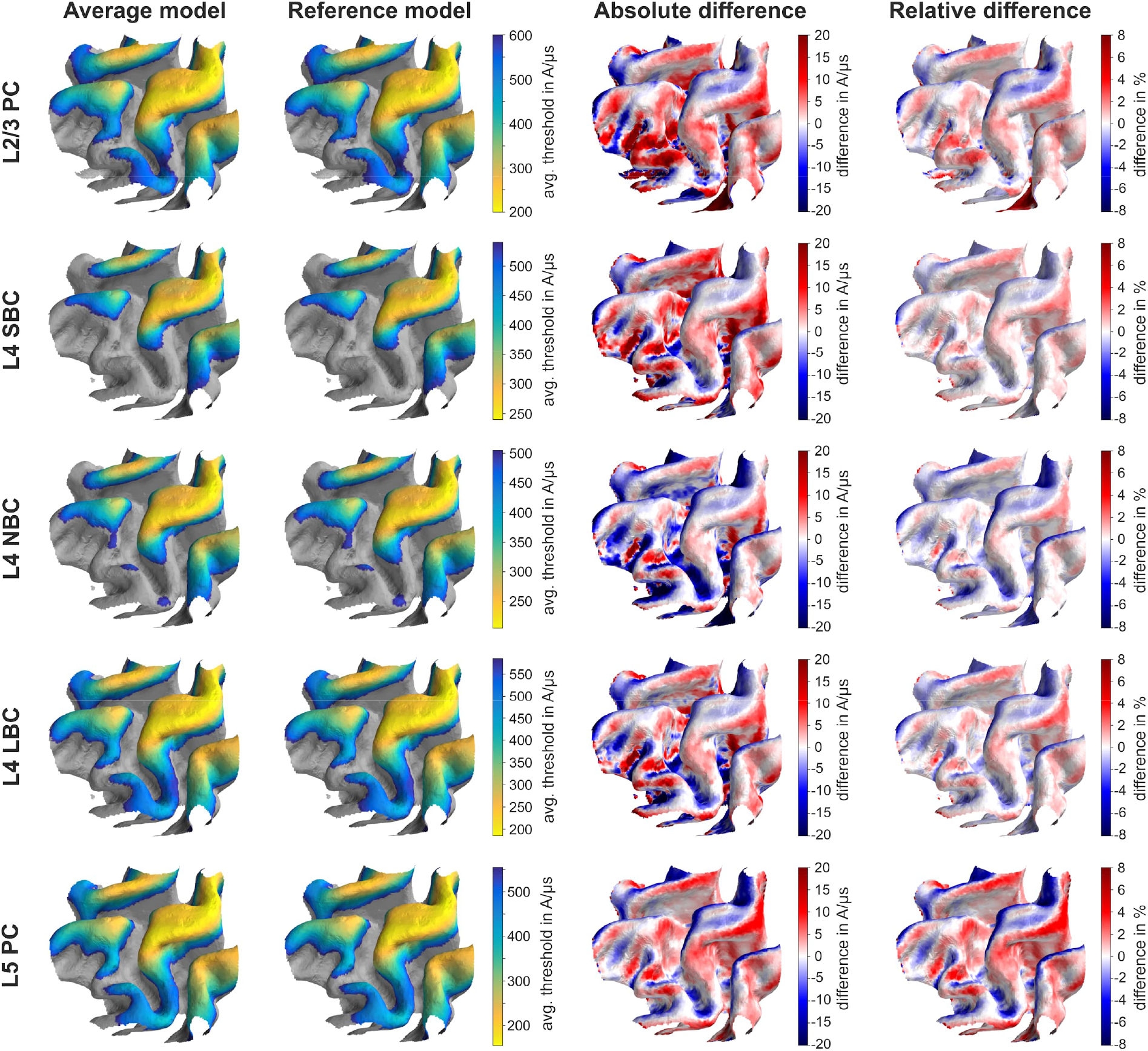
Comparison of stimulation intensity threshold maps (in A/µs) for monophasic excitation determined using the average model and the reference model: The first two rows show the stimulation threshold maps (in A/µs) of the L2/3 PC and the last two rows of the L5 PC between the average model (first column) and the reference model (second column). The last two columns show the absolute and relative difference between the models. It is assumed that the TMS coil is located over the M1 area with an orientation of 45° towards the *fissura longitudinalis*. The maps indicate the stimulation strength of the TMS device in A/µs, which is required to stimulate this cortical area for this particular coil position and orientation. The underlying electric field distribution and field direction is shown in Fig. 4. The results for biphasic excitation are shown in Figure S9.

To quantify the differences between the models, we determined the normalized root mean square error (NRMSE):

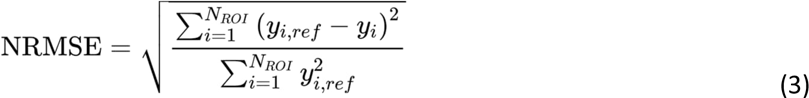

where *y_i, ref_* denotes the thresholds of the reference simulations in the *i*-th ROI element and *y_i_* the thresholds from the average model. Additionally, we calculated the mean absolute percentage error (MAPE) quantifying the prediction accuracy of the average models:

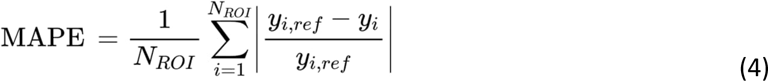

and the coefficient of determination (R^2^) quantifying the proportion of the total variance explained by the average model:

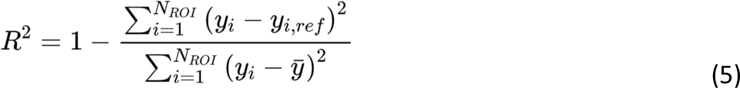

where is the mean of the average threshold model.

The histograms of the relative differences are shown in Fig. 14 together with the different error measures. The distribution of relative differences is relatively symmetric and the means are close to zero. The remaining variance results from the inhomogeneity of the electric field across the neurons. Systematic field distortions in a particular direction across neurons, such as those occurring at the gyral crowns, are neglected and result in deviations from the exact reference model because in the simplified model, only the decay of the electric field across the somatodendritic axis can be accounted for due to averaging over the azimuthal angle.

**Figure 14:**
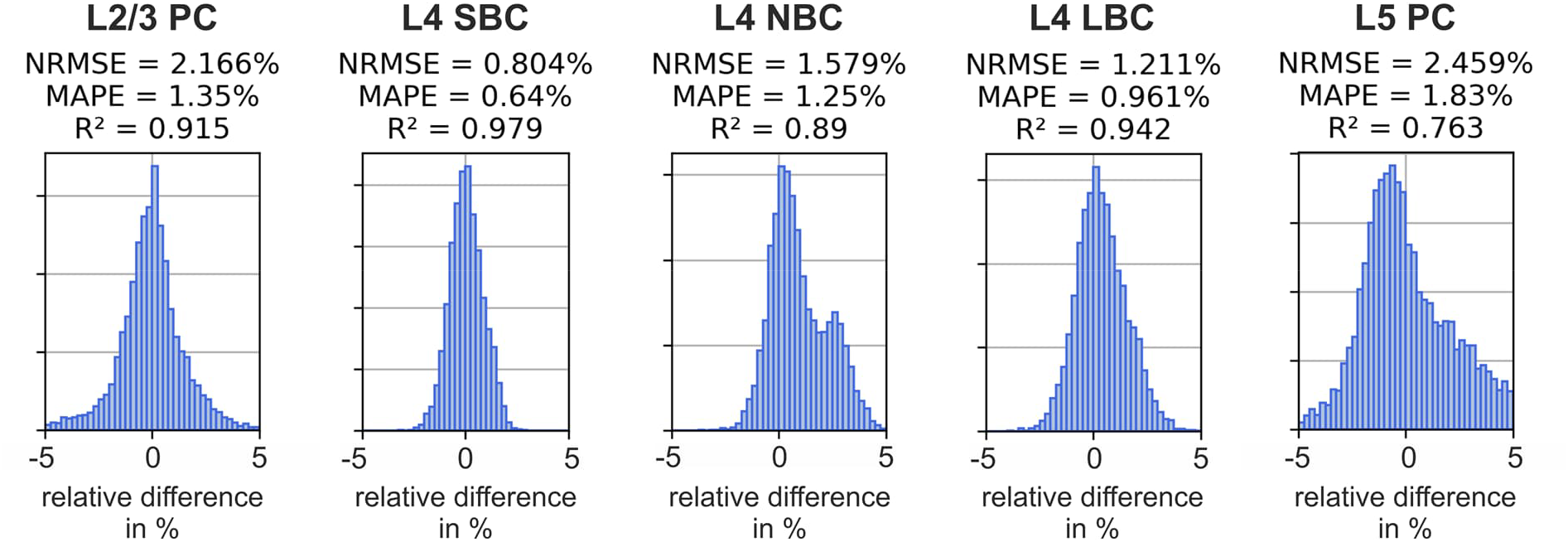
Differences of the threshold maps between the average model and the reference model for monophasic excitation. Histograms of the relative difference between the reference model and the average threshold model over the ROI elements. Normalized root mean square error (NRMSE), mean absolute percentage error (MAPE), and coefficient of determination (R^2^) for L2/3 PC and L5 PC with monophasic and biphasic excitation. The results for biphasic excitation are shown in Fig. S10.

### 3.7 Validation

The predicted orientation sensitivity of the neurons is compared to the orientation sensitivity of MEPs. The polar plot in Fig. 15 shows the MEP amplitudes for different electric field angles and stimulation intensities together with the predictions of the recruitment rate from the average threshold model from L5 PC under monophasic excitation. To make both representations comparable, the data were normalized to their respective maximum values. The average threshold model closely matches the orientation sensitivity of MEPs and the NRMSE between the experimental data and model predictions is 8.5%. It can be clearly observed that the directional sensitivity is more pronounced for low stimulation intensities close to rMT than for higher ones. The cortical column cosine model resembles the general behavior of the directional sensitivity of the MEPs at low stimulation intensities, but cannot represent the stimulation behavior in the transition to higher stimulation intensities. The ball-and-stick model was also not able to reflect the direction sensitivity over the investigated parameter range for both the incident angle *θ* and the stimulation intensity.

**Figure 15:**
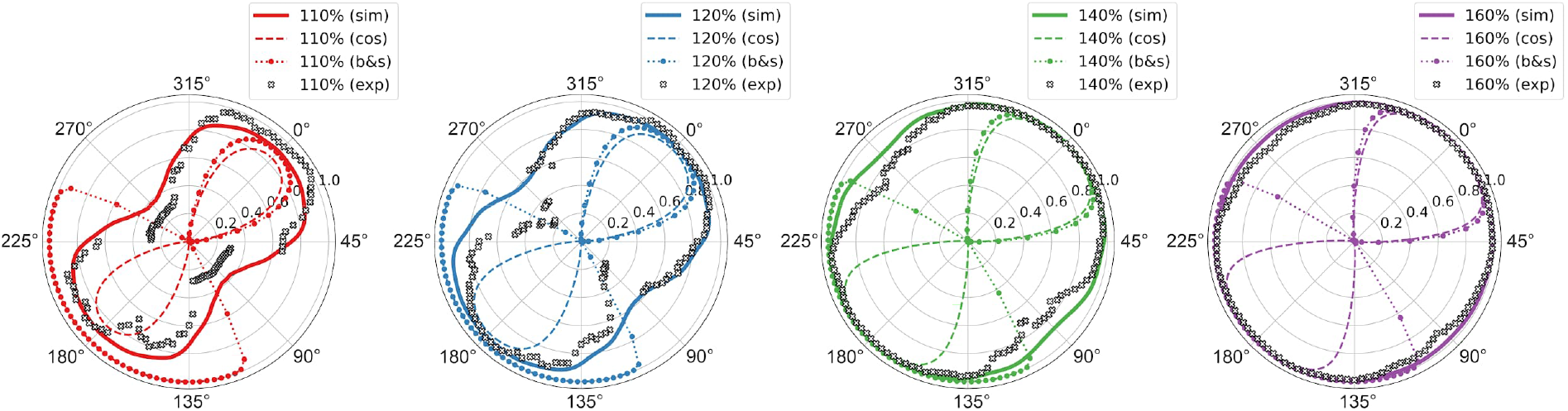
Comparison between directional sensitivity of motor evoked potentials and the derived recruitment rate from the theoretical neuronal response model: The plots show the directional sensitivity of measured MEPs as black crosses (exp) at different stimulation intensities with respect to resting motor threshold of subject 16 from Souza et al. (2022). The solid lines (sim) show the corresponding trajectory of the recruitment rate along *θ* assuming a constant electric field along the somato-dendritic axis (Δ|**Ẽ**|=0 %/mm), and the dashed and the dotted lines show the predictions of the directional sensitivity of the MEPs according to the cortical column cosine model (cos) and the ball-and-stick model (b&s). The MEPs were normalized to their maximum values for comparability. The NRMSE between the experimental data and the recruitment rate model (sim) is 8.5%.

## 4 Discussion

In order to link the predicted electric field to actual neural activation, a range of different proposals with varying degrees of complexity have been put forward. The simplest approach just considers the magnitude of the electric field as a proxy for the activation strength (e.g., Weise et al., 2021), without any dependency on direction or local gradient of the field. This method disregards experimental observations and theoretical considerations showing that the activation threshold does indeed depend on the incidence angle between the field direction and the orientation of the axons (Rushton, 1927; Rudin and Eisenman, 1954; Ranck, 1975). This consideration led to the proposal of the cortical column cosine model (Fox et al., 2004), which is based on the assumption that axons aligned with the somato-dendritic axis (i.e., perpendicular to the cortical surface) dominate the stimulation process, and therefore predicts that only the projection of the electric field onto that axis has an effect. As a consequence, purely tangential fields would lead to no stimulation.

However, it is known that the axonal arbors of cortical cells are much branched and cover all directions (Aberra et al., 2020). In line with this, in earlier TMS motor mapping experiments (Weise et al., 2020, Numssen et al., 2021, Weise et al., 2023a), we could show that the tangential field component does indeed have a substantial predictive power towards the resulting motor evoked potentials. In fact, it was even considerably more powerful than the radial component (i.e., the one aligned with the somato-dendritic axis), which can be understood in the light of the cortical geometry: At the gyral crowns, the field is largely tangential to the cortical surface (thus, having less impact), but its magnitude is much larger due to the greater proximity to the coil, thus overcompensating the former effect. In contrast, at the sulcal walls, located more distant to the coil, the field is radial, but much weaker, and therefore often does not effectively stimulate.

In order to obtain a more accurate account of the coupling between the electric field and the activation states of cortical neurons, detailed biological models based on realistic neuronal geometry and realistic Hodgkin-Huxley-like neural dynamics have been proposed (Aberra et al., 2018, 2020). These models have the potential to deliver a detailed and accurate picture of neuronal activation by TMS. However, they are computationally extremely demanding and therefore hardly suitable for routine applications, such as mapping or dosing. Moreover, the utilized neural geometries must be considered as samples of a distribution and do not account for any precise individual cortical architecture. This suggests that the predictions of these models should be representable in low-parametric models without much loss.

In our study, we attempt to bridge the gap between, on the one hand, the imprecise oversimplification of the magnitude, cortical column cosine, and ball-and-stick models and, on the other hand, the unwieldy and time-consuming biologically realistic modeling. The model we propose is as easily applicable as the former, while it very closely approximates the predictions of the latter. It determines the stimulation thresholds as functions of field angle with respect to the somato-dendritic axis, intensity, pulse waveform, and field decay along the somato-dendritic axis, and only requires the induced electric field as an input variable. Comparison with reference simulations with a detailed neuronal model yielded normalized root mean square errors of only 1.5-2.5%. It should be emphasized, however, that our model is not independent, but depends, for initial calibration, on a biologically realistic model based on the principal approach of Aberra and colleagues (2018, 2020).

Our model predicts a certain dependence of the stimulation threshold on the angle of incidence of the electric field, which is more pronounced for monophasic pulses than for biphasic pulses. However, compared to the predictions from simplified models such as the cortical column cosine model or ball-and-stick neurons, this dependence is much more moderate. This allows tangential electric fields of sufficient strength to contribute to the stimulation, as has been observed in previous experimental studies (Weise et al., 2020, Numssen et al., 2021). In particular, the L2/3 PC require 108%, L4 SBC require 113%, L4 NBC require 110%, L4 LBC require 114%, and L5 PC require 120% of the longitudinal stimulation strength (*θ*=0°) at *θ*=90° for monophasic excitation, respectively. For a comparison of the directional sensitivity profiles of our model and the cortical column cosine and ball-and-stick models, see Fig. 5-9.

These findings are confirmed by a comparison to the experimentally observed orientation sensitivity of MEPs (Fig. 14), where a difference of only 8.5% was observed between model predictions and experimental data. We observed a high directional sensitivity at low stimulation intensities close to the motor threshold, while at higher stimulation intensities the directional sensitivity rapidly decreases. While our model appears to be a quite good predictor of the directional sensitivity observed by Souza et al. (2022), there are also deviations. This is mainly explainable in the light of some important discrepancies between the assumptions underlying our model and those made by these authors. First, the results of Souza and colleagues are based on electric fields predicted using a spherical head model, while our model works with a realistic head model. Second, Souza’s report is based on the electric field direction with respect to the global coordinate system, while our angle definition is local and changes across the strongly curved cortical surface. Third, the location of the neuronal populations that mediate the relationship between stimulation and MEP is only roughly known in Souza et al. (2022). It may therefore be that the field angles at that location are different from those predicted for the assumed target spot. Accordingly, for an even better comparison, the currents in the multi-coil array would have to be optimized subject- and target-specifically to realize an ideal rotation of the electric field at a constant field strength in the target. This in turn requires precise knowledge of the target and thus a prior mapping of the motor cortex such as in Weise et al. (2023a).

The major advantage of the presented model is its simplicity without sacrifice of realism. The availability of look-up tables of threshold maps and recruitment rates allows for the simple construction of interpolators and functions for computation. Alternatively, polynomial-based surrogate models based on generalized polynomial chaos (gPC; Weise et al., 2020b) can be used for this purpose and provide high accuracy. Examples are given in the repository of Weise et al. (2023b).

Importantly, the model is easy to adapt and refine, if more or better information about the neuronal geometry of particular tissues becomes available, using the provided scripts and simulation code (https://github.com/TorgeW/TMS-Neuro-Sim). Already in this study, we were able to distinguish between the stimulation thresholds and distributions among various distinct cell types. We observed that L5 PCs had the lowest thresholds compared to all other cell types studied, followed by L4 LBC and L2/3 PCs, which had 10% and 22% higher thresholds, respectively. This “library” of cellular stimulation profiles may be extended in the future. By comparison with experimentally observed stimulation profiles, such cell-specific sensitivity profiles may potentially allow for testing hypotheses about which cells are actually stimulated in particular experimental situations.

These traits allow for efficient implementation and extension of TMS models in the context of optimization, mapping, and dosing without the need to implement time consuming and complicated neuron models. Especially in the field of cognitive TMS experiments, where an adequate dosing strategy is still subject to research, the gained knowledge could significantly contribute to the identification of the effectively stimulated regions but also to exclude regions that are not eligible for stimulation due to the underlying electric field distribution and orientation relative to the cortex.

The threshold maps have revealed interesting parameter combinations of *θ* and Δ|**Ẽ**| that enable particularly effective stimulation. Here, an interesting observation is that an increase of the electric field along the somato-dendritic axis of the neurons (Δ|**Ẽ**|>0) from the GM surface to the WM surface is as likely as a decrease and is usually in the range of ±20%/mm (Fig. 3b). Future optimization studies could be directed towards identifying coil positions and orientations that realize these parameter combinations in the targeted region. As a result, such new optimization strategies would have great potential to significantly enhance the overall efficacy of TMS and reduce the required dose. At the same time, the optimization criterion can be extended such that the electric field is oriented to *prevent* stimulation of other brain regions by targeting particularly high thresholds. This principled approach of multi-objective optimization was also taken up by Lueckel et al. (2022) in the framework of electric field and connectivity optimized TMS targeting.

Another area of application for the presented models is in the extension of existing mapping procedures (Weise et al., 2020, Numssen et al., 2021; Weise et al., 2023a), as mentioned previously. Instead of the electric field magnitude, some kind of effective electric field could be used as a regressor for localization. It is noted that the integration of the stimulation thresholds into the analysis procedures occurs solely at the modeling level, thus improving the efficiency of the mapping procedures without increasing the experimental effort. Stronger even, the fact that we have an estimate of the stimulation threshold at every cortical location, we can successively exclude locations which are stimulated without an observable effect (e.g., MEP), and thereby even decreasing the experimental effort.

### Limitations of the study

The number of L2/3 and L5 neurons available was limited. We were able to significantly expand the original dataset of Aberra et al. (2018) and Aberra et al. (2020), but especially for the calculation of the recruitment rate, a higher number of neurons would increase the model accuracy. This is especially true for incident angles where threshold variances are high.

Moreover, we limited the analysis to pyramidal cells in L2/3, L5, and Basket cells in L4, which take a major part in generalized cortical circuits (Di Lazzaro et al., 2012). However, it is known that other cell types like spiny stellates in L4, also play a major role in the stimulation of cortical microcircuits. The development of average threshold models for other cell types is straightforward using the tools provided in the repository Weise et al. (2023b) and the Python package *TMS-Neuro-Sim* (https://github.com/TorgeW/TMS-Neuro-Sim) if the appropriate morphologies and parameterizations are available.

In the modeling, we also neglected the effect of the presence of the neurons and other cells to the external electric field. While for the macroscopic field estimation, these structures are already accounted for through the (macroscopically acquired) tissue conductivity, at the microscale, the presence of low conducting membranes might cause local deviations from that macroscopically predicted field, which may have an effect on the actual stimulation of neurons.

### Future work

Insights into the stimulation behavior of neurons are essential to develop realistic coupling models for downstream neuronal mass models along the lines of Montbrió et al. (2015) or Jansen and Rit (1995), which in turn could be used to model the dynamic processes of entire populations of neurons, such as the D- and I-wave dynamics in the motor cortex (Di Lazzaro et al. 1998,; Di Lazzaro et al., 2012; Ziemann, 2020).

In further follow-up studies, the degree to which the spatial fine-structure of the electric field is affected by the high membrane resistance of the neurons should be investigated. The resulting change in the electric field distribution may have a non-negligible influence on the local electric field angles and magnitudes, which in turn change the stimulation thresholds. However, this will require very detailed volume conductor models of whole cortical columns or at least geometric information about the neuron surfaces and their position with respect to each other. It is expected that this type of model will require high computational power to solve and is far from being routinely used in daily TMS experiments and that it will lead to an anisotropic macroscopic conductivity profile as well as potentially modified sensitivity profile due to local electric field fluctuations. The former can be estimated with diffusion-weighted MRI (Güllmar et al., 2010), but its influence on the stimulation behavior on a micro- and mesoscopic scale is yet unknown. The goal of such a study could be the derivation of a new generation of low-parametric models, in a similar sense as in this study, in order to be able to apply the gained knowledge in practice.

A further step towards a more accurate biophysical modeling of the stimulation processes may consist in the consideration of the back reaction of the neurons to the extracellular potential when action potentials are generated. Active ion transport alters the total electric field and can lead to mutual interference (cross-talk) between neurons. For such a model, the neurons can no longer be considered separately, but must be simulated as a unit in the form of a cortical column or similar. Such a model approach can be combined with the previous one, but it is expected that the required computing time will be even higher to solve it.

## Acknowledgements

This work was partially supported by the German Science Foundation (DFG) (grant number WE 59851/2 to K.W. KN 588/10-1 to T.R.K.), Lundbeckfonden (grant no. R244-2017-196 and R313-2019-622) and the NVIDIA Corporation (donation of one Titan Xp graphics card to K.W.). V.H.S. is funded by the Academy of Finland (decision No 349985). A.J. and W.V.G are supported by funding to the Blue Brain Project, a research center of the École polytechnique fédérale de Lausanne (EPFL), from the Swiss government’s ETH Board of the Swiss Federal Institutes of Technology.

## Conflicts of interest

The authors declare that they have no known competing financial interests or personal relationships that could have appeared to influence the work reported in this paper.

## Notes

https://osf.io/c8j35/

### Competing Interest Statement

The authors have declared no competing interest.

