## Supplemental Material for "Directional Sensitivity of Cortical Neurons Towards TMS Induced Electric Fields"

Konstantin Weise[1,2*+], Torge Worbs[1,3+], Benjamin Kalloch[1,4], Victor H. Souza[5], Aurélien Tristan Jaquier[6], Werner Van Geit[6], Axel Thielscher[3,7], Thomas R. Knösche[1,4]

[1]Methods and Development Group “Brain Networks”, Max Planck Institute for Human Cognitive and Brain Sciences, Stephanstr. 1a, 04103 Leipzig, Germany.

[2]Department of Clinical Medicine, Aarhus University, DNK-8200, Aarhus, Denmark

[3]Technical University of Denmark, Magnetic Resonance Section, Department of Health Technology, Kongens Lyngby, Denmark.

[4]Technische Universität Ilmenau, Institute of Biomedical Engineering and Informatics, Gustav-Kirchhoff-Straße 2, 98693 Ilmenau, Germany.

[5]Department of Neuroscience and Biomedical Engineering, Aalto University School of Science, Espoo, Finland

[6]Blue Brain Project, École polytechnique fédérale de Lausanne (EPFL), Biotech Campus, 1202 Geneva, Switzerland

[7]Danish Research Centre for Magnetic Resonance, Section for Functional and Diagnostic Imaging and Research, Copenhagen University Hospital Amager and Hvidovre, Denmark.

* CORRESPONDING AUTHOR

+ contributed equally

Konstantin Weise; Max Planck Institute for Human Cognitive and Brain Sciences, Stephanstr. 1a, 04103 Leipzig, Germany; Technische Universität Ilmenau, Advanced Electromagnetics Group, Helmholtzplatz 2, 98693 Ilmenau, Germany;, phone: +49 341 9940-2580

### Stimulation behavior of L2/3 PCs for biphasic excitation

The results for L2/3 PCs when excited with biphasic TMS pulses are shown in Fig. S1. The threshold map in dependence of $\vartheta$ and $\Delta|\tilde{\mathbf{E}}|$ is shown in Fig. S1a, a slice of the threshold map together with the individual neuron thresholds are shown in Fig. S1b for $\Delta|\tilde{\mathbf{E}}|$=0 %/mm, and the recruitment rate is shown in Fig. S1c. Again, the lowest thresholds can be observed when the electric field is parallel to the somato-dendritic axis, i.e. for $\vartheta$=0° and $\vartheta$=180°. The difference between both stimulation conditions is lower compared to monophasic pulses (Fig. 5) due to the existence of both field directions in case of a biphasic excitation. The thresholds for tangential electric fields ($\vartheta$=90°) are about 11% higher compared to normal electric fields ($\vartheta$=0° and $\vartheta$=180°). The lowest thresholds can be observed for $\vartheta$=0° in combination with a positive electric field change along the somato-dendritic axis at $\Delta|\tilde{\mathbf{E}}|$=30 %/mm. Directional sensitivity of the L2/3 PCs is clearly observed, but not as pronounced as with monophasic pulses and in general, only about 85% of the electric field strength is needed to reach the stimulation threshold compared to monophasic pulses.

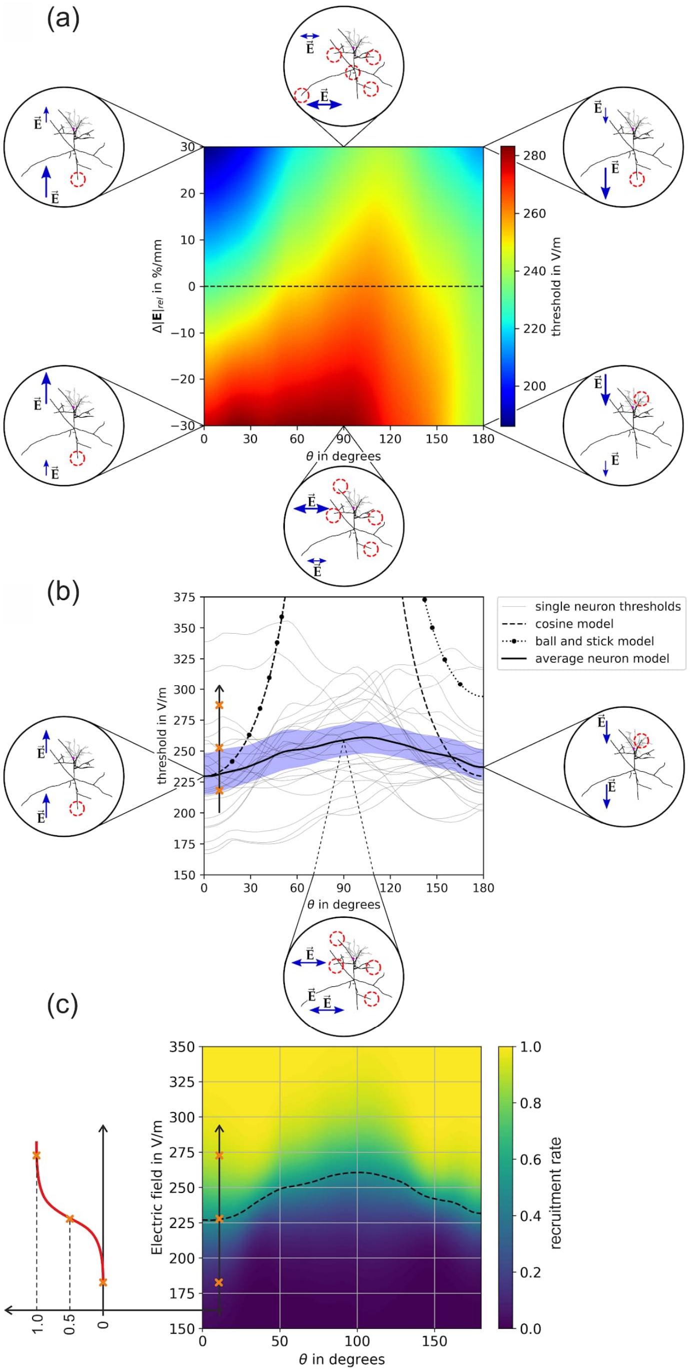


Figure S1: Stimulation behavior of L2/3 PCs for biphasic excitation: (a) Threshold map in dependence of the polar angle $\vartheta$ and the relative change of the electric field over the somato-dendritic axis $\Delta|\tilde{\mathbf{E}}|$. The insets show the locations of excitation, the red circles indicate the activated terminals. Blue arrows indicate the electric field direction and magnitude; (b) Thresholds of individual neurons for $\Delta|\tilde{\mathbf{E}}|$=0 %/mm along the dashed line in (a). The blue area shows the 95th percentile of the confidence interval of the mean. The equivalent cortical column cosine model is $y(\theta) = \hat{y}|\cos(\theta)|^{-1}$ with $\hat{y}$=229.71 V/m (dashed line); the axon parameters of the equivalent ball and stick model are $l = 200\,\mu\mathrm{m}$ and $d = 6.4\,\mu\mathrm{m}$ (dotted line); (c) Recruitment rate for $\Delta|\tilde{\mathbf{E}}|$=0 %/mm derived from the individual neuron activation in (b) by integrating over the electric field thresholds. The dashed line indicates the electric field intensity where the recruitment rate is 0.5.

### Stimulation behavior of L4 SBCs for biphasic excitation

The results of the average response model of L4 SBCs in case of a biphasic excitation is shown in Fig. S2. A pronounced directional sensitivity can also be observed for this cell type. Again, lowest thresholds can be observed when the electric field is parallel to the somato-dendritic axis ($\vartheta$=0° and $\vartheta$=180°). The thresholds are about 10% higher when the external electric field is tangential to the cells ($\vartheta$=90°). The thresholds are slightly affected if the electric field changes along the somato-dendritic axis ($\Delta|\tilde{\mathbf{E}}|\neq 0$ %/mm). Compared to other cells, the average threshold is about 16% and 45% higher for L4 SBCs than for L2/3 PCs and L5 PCs, respectively.

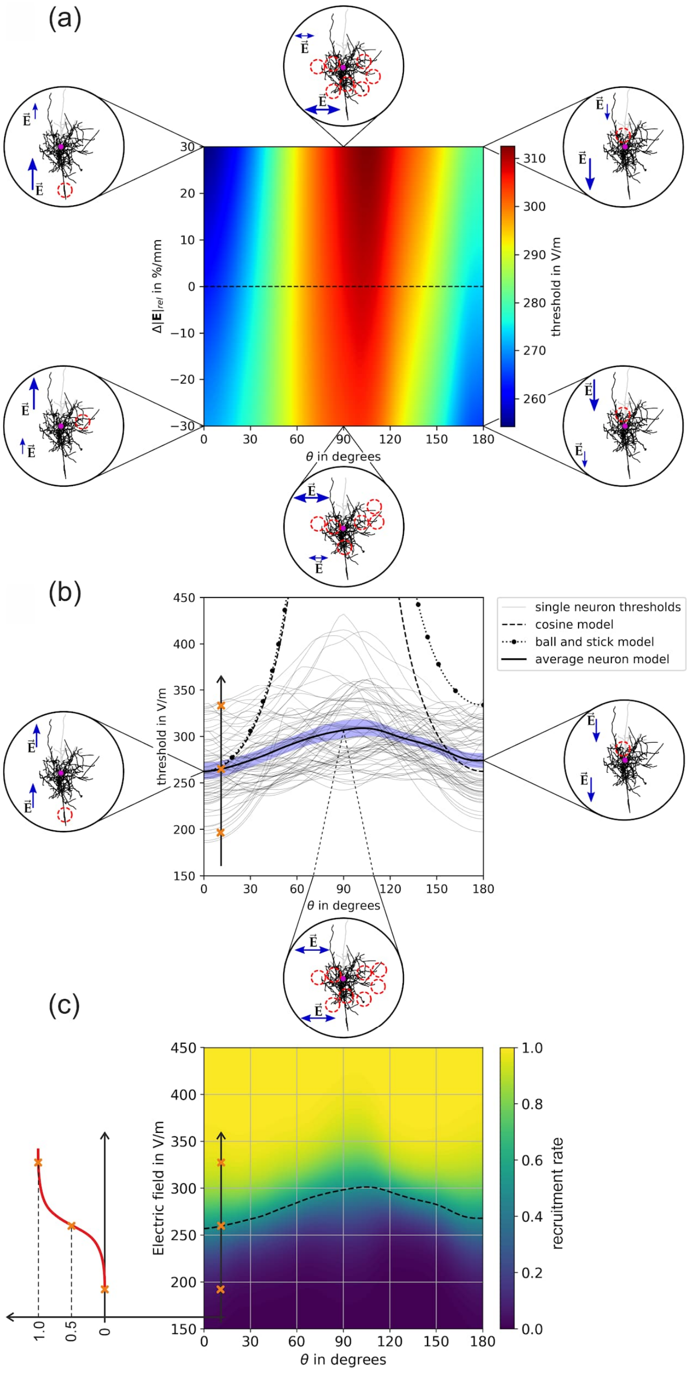


Figure S2: Stimulation behavior of L4 SBCs for biphasic excitation: (a) Threshold map in dependence of the polar angle $\vartheta$ and the relative change of the electric field over the somato-dendritic axis $\Delta|\tilde{\mathbf{E}}|$. The insets show the locations of excitation, the red circles indicate the activated terminals. Blue arrows indicate the electric field direction and magnitude; (b) Thresholds of individual neurons for $\Delta|\tilde{\mathbf{E}}|$=0 %/mm along the dashed line in (a). The blue area shows the 95th percentile of the confidence interval of the mean. The equivalent cortical column cosine model is $y(\theta) = \hat{y}|\cos(\theta)|^{-1}$ with ŷ=262.33 V/m (dashed line); the axon parameters of the equivalent ball and stick model are $l = 150\,\mu\mathrm{m}$ and $d = 8\,\mu\mathrm{m}$ (dotted line); (c) Recruitment rate for $\Delta|\tilde{\mathbf{E}}|$=0 %/mm derived from the individual neuron activation in (b) by integrating over the electric field thresholds. The dashed line indicates the electric field intensity where the recruitment rate is 0.5.

### Stimulation behavior of L4 NBCs for biphasic excitation

The results of the average response model of L4 NBCs in case of a monophasic excitation is shown in Fig. S3. Their axonal arborization is distinct from pyramidal cells because they form intricate networks of branches that wrap around the soma of nearby pyramidal cells, forming a characteristic "basket" structure. Their axonal structure is generally more isotropic compared to pyramidal cells or SBCs and LBCs. This also affects the stimulation properties and explains the weaker directional sensitivity of these cells observed in Fig. S3a and b. The thresholds for tangential electric fields are about 11% higher compared to normal electric fields ($\vartheta$=0° and $\vartheta$=180°). On average, the thresholds of L4 NBCs are 2% lower to L2/3 PC and 22% higher compared to L5 PCs, respectively.

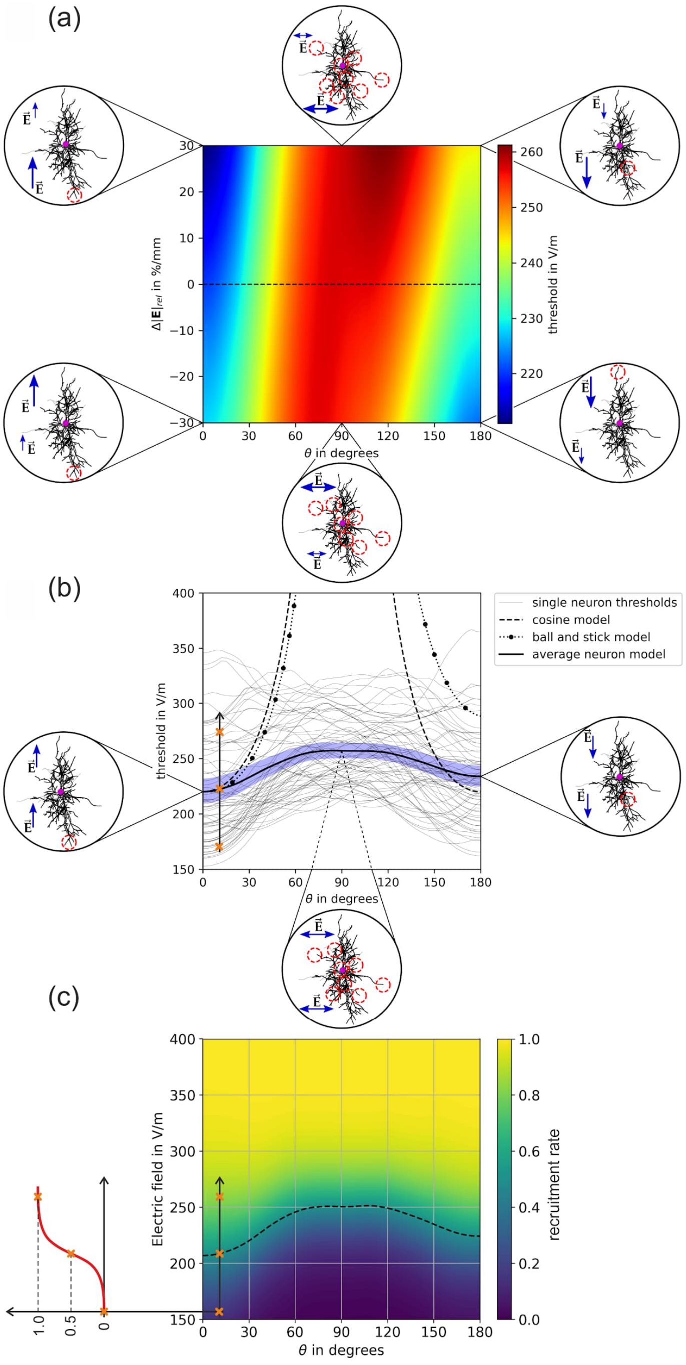


Figure S3: Stimulation behavior of L4 NBCs for biphasic excitation: (a) Threshold map in dependence of the polar angle $\vartheta$ and the relative change of the electric field over the somato-dendritic axis $\Delta|\tilde{\mathbf{E}}|$. The insets show the locations of excitation, the red circles indicate the activated terminals. Blue arrows indicate the electric field direction and magnitude; (b) Thresholds of individual neurons for $\Delta|\tilde{\mathbf{E}}|$=0 %/mm along the dashed line in (a). The blue area shows the 95th percentile of the confidence interval of the mean. The equivalent cortical column cosine model is $y(\theta) = \hat{y}|\cos(\theta)|^{-1}$ with ŷ=220.05 V/m (dashed line); the axon parameters of the equivalent ball and stick model are $l = 200\,\mu\mathrm{m}$ and $d = 4.5\,\mu\mathrm{m}$ (dotted line); (c) Recruitment rate for $\Delta|\tilde{\mathbf{E}}|$=0 %/mm derived from the individual neuron activation in (b) by integrating over the electric field thresholds. The dashed line indicates the electric field intensity where the recruitment rate is 0.5.

### Stimulation behavior of L4 LBCs for biphasic excitation

The threshold results of L4 LBCs for monophasic excitation are shown in Fig. S4. Compared to PCs, LBCs exhibit a high degree of collateralization in their axonal tree. They can have multiple branches and collaterals that extend in different directions within the same cortical layer or across layers. A distinct directional sensitivity of the thresholds can be again observed together with an asymmetric modulation when the electric field changes along the somato-dendritic axis. The thresholds for tangential electric fields are about 8% higher compared to normal electric fields ($\vartheta$=0° and $\vartheta$=180°). On average, the thresholds of L4 LBCs are 11% lower than L2/3 PC and 10% higher compared to L5 PCs, respectively. Of all the basket cells investigated, the LBCs have the lowest thresholds. The average thresholds for LBCs are about 23% and 10% lower compared to SBCs and NBCs, respectively.

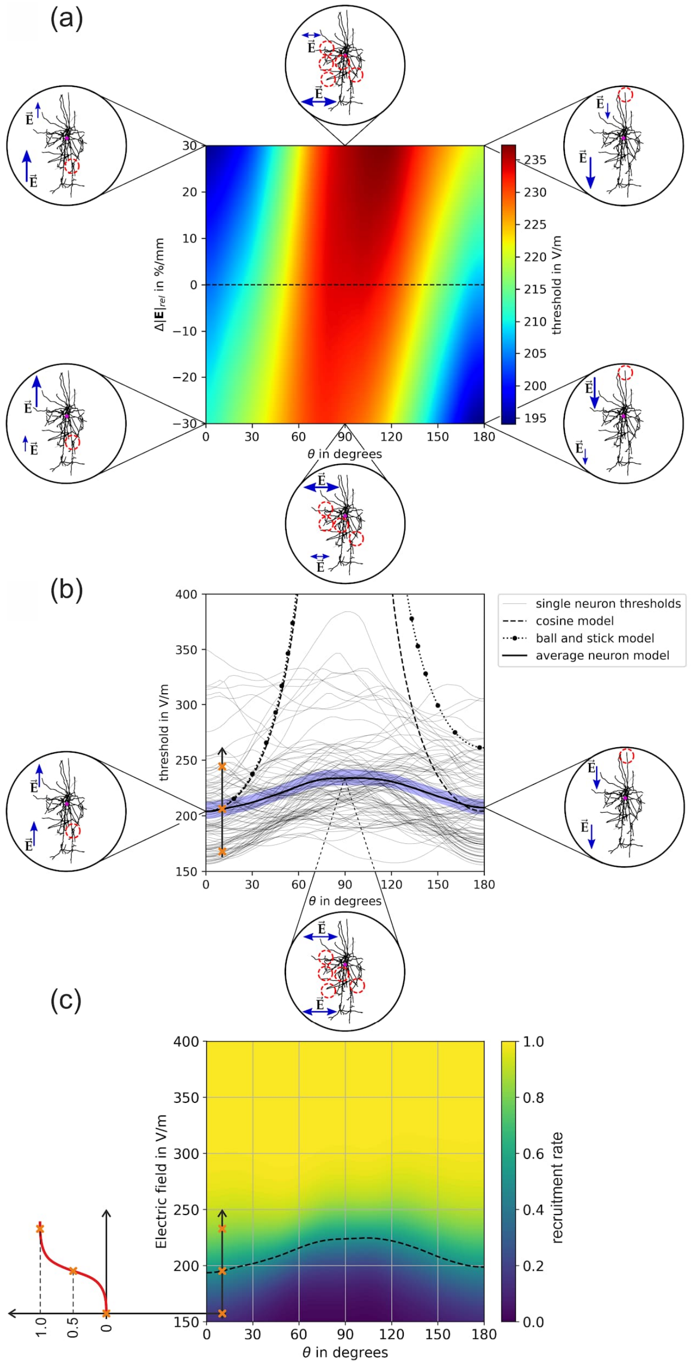


Figure S4: Stimulation behavior of L4 LBCs for biphasic excitation: (a) Threshold map in dependence of the polar angle $\vartheta$ and the relative change of the electric field over the somato-dendritic axis $\Delta|\tilde{\mathbf{E}}|$. The insets show the locations of excitation, the red circles indicate the activated terminals. Blue arrows indicate the electric field direction and magnitude; (b) Thresholds of individual neurons for $\Delta|\tilde{\mathbf{E}}|$=0 %/mm along the dashed line in (a). The blue area shows the 95th percentile of the confidence interval of the mean. The equivalent cortical column cosine model is $y(\theta) = \hat{y}|\cos(\theta)|^{-1}$ with $\hat{y}$=204.06 V/m (dashed line); the axon parameters of the equivalent ball and stick model are $l = 250\,\mu\mathrm{m}$ and $d = 5.6\,\mu\mathrm{m}$ (dotted line); (c) Recruitment rate for $\Delta|\tilde{\mathbf{E}}|$=0 %/mm derived from the individual neuron activation in (b) by integrating over the electric field thresholds. The dashed line indicates the electric field intensity where the recruitment rate is 0.5.

### Stimulation behavior of L5 PCs for biphasic excitation

The results for biphasic excitation of L5 PCs is shown in Fig. S5. The profile of the threshold map in Fig. S5a resembles the monophasic case. It can be observed that the variance of the stimulation thresholds between the cells in Fig. S5b is lower across the polar angle $\vartheta$ compared to monophasic excitation (Fig. 9b). The thresholds for tangential electric fields ($\vartheta$=90°) are about 16% higher compared to normal electric fields ($\vartheta$=0° and $\vartheta$=180°). The relative electric field change required to stimulate L5 PCs most efficiently is reversed compared to the monophasic case. The most stimulation of L5 PCs can be achieved with electric fields with a polar angle of $\vartheta$=0° and a positive relative electric field change ($\Delta|\tilde{\mathbf{E}}|>0$) across the somato-dendritic axis or with an angle of $\vartheta$=180° together with a negative field decay ($\Delta|\tilde{\mathbf{E}}|<0$). Likewise, the stimulation locations for $\vartheta$=0° and $\vartheta$=180° are also reversed compared to monophasic stimulations. For biphasic stimulations, axon collaterals in the upper part of the cell are stimulated for $\vartheta$=180°, which may connect to other cell populations within the cortex. In case of antidromic electric fields at $\vartheta$=0°, lower parts of the axons are stimulated, indicating the activation of cortico-spinal connections.

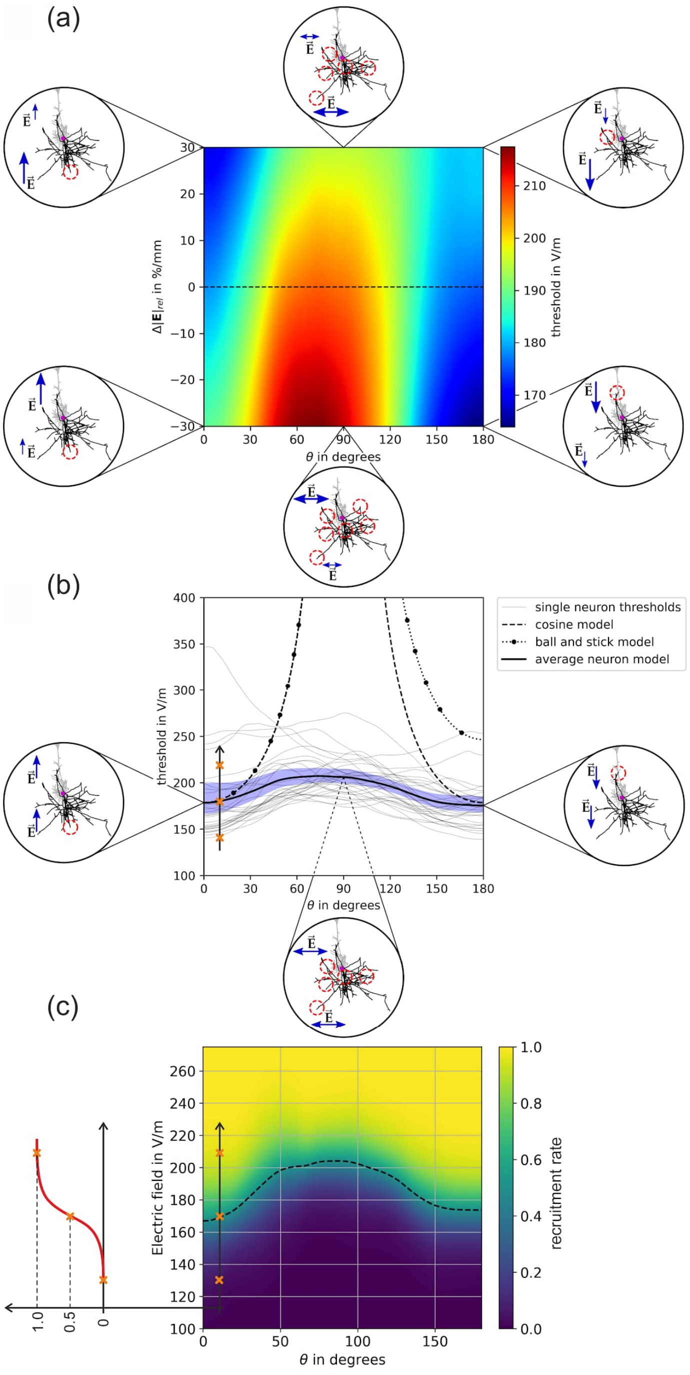


Figure S5: Stimulation behavior of L5 PC for biphasic excitation: (a) Threshold map in dependence of the polar angle $\vartheta$ and the relative change of the electric field over the somato-dendritic axis $\Delta|\tilde{\mathbf{E}}|$. The insets show the locations of excitation, the red circles indicate the activated terminals. Blue arrows indicate the electric field direction and magnitude; (b) Thresholds of individual neurons for $\Delta|\tilde{\mathbf{E}}|$=0 %/mm along the dashed line in (a). The blue area shows the 95th percentile of the confidence interval of the mean. The equivalent cortical column cosine model is $y(\theta) = \hat{y}|\cos(\theta)|^{-1}$ with $\hat{y}$=178.43 V/m (dashed line); the axon parameters of the equivalent ball and stick model are $l = 200\,\mu\text{m}$ and $d = 3.8\,\mu\text{m}$ (dotted line); (c) Recruitment rate for $\Delta|\tilde{\mathbf{E}}|$=0 %/mm derived from the individual neuron activation in (b) by integrating over the electric field thresholds. The dashed line indicates the electric field intensity where the recruitment rate is 0.5.

### Recruitment order and relative threshold ranges for biphasic excitation

Similar to monophasic excitations, L5 PCs have the lowest thresholds compared to all other investigated cell types. The L4 LBCs have the second lowest thresholds followed by the L2/3 PCs and the L4 NBCs. The small basket cells are again stimulated only at higher stimulation intensities.

In particular, the L2/3 PC require 113%, L4 SBC require 117%, L4 NBC require 117%, L4 LBC require 115%, and L5 PC require 115% of the longitudinal stimulation strength ($\vartheta$=0°) at $\vartheta$=90° for biphasic excitation, respectively.

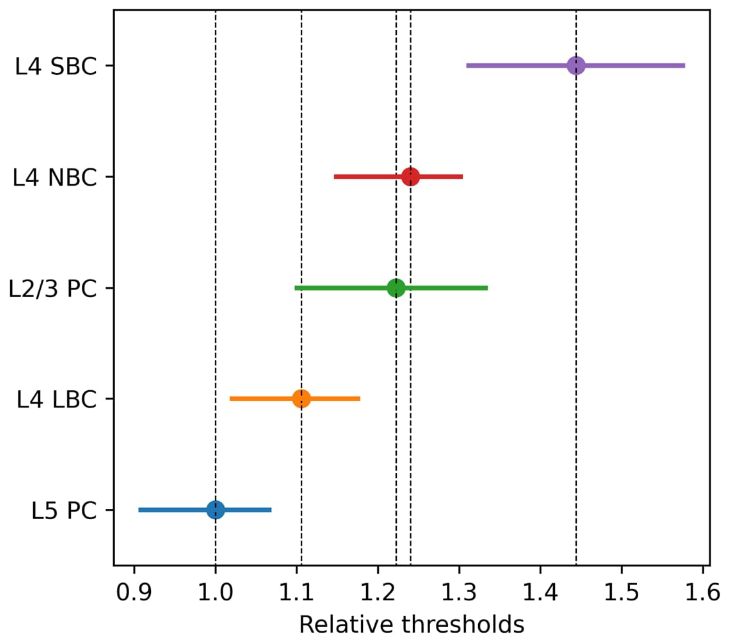


Figure S6: Recruitment order and relative threshold ranges of pyramidal and basket cells for monophasic TMS excitation: Threshold ranges of all investigated cell types relative to the mean of L5 PCs, is shown assuming a constant electric field along the somatodendritic axis ($\Delta|\tilde{E}|$=0 %/mm). The dots indicate the mean thresholds and the ranges stem from the variability across the polar angle $\vartheta$ from 0° to 180°.

### Sensitivity analysis for L5 PCs under biphasic excitation

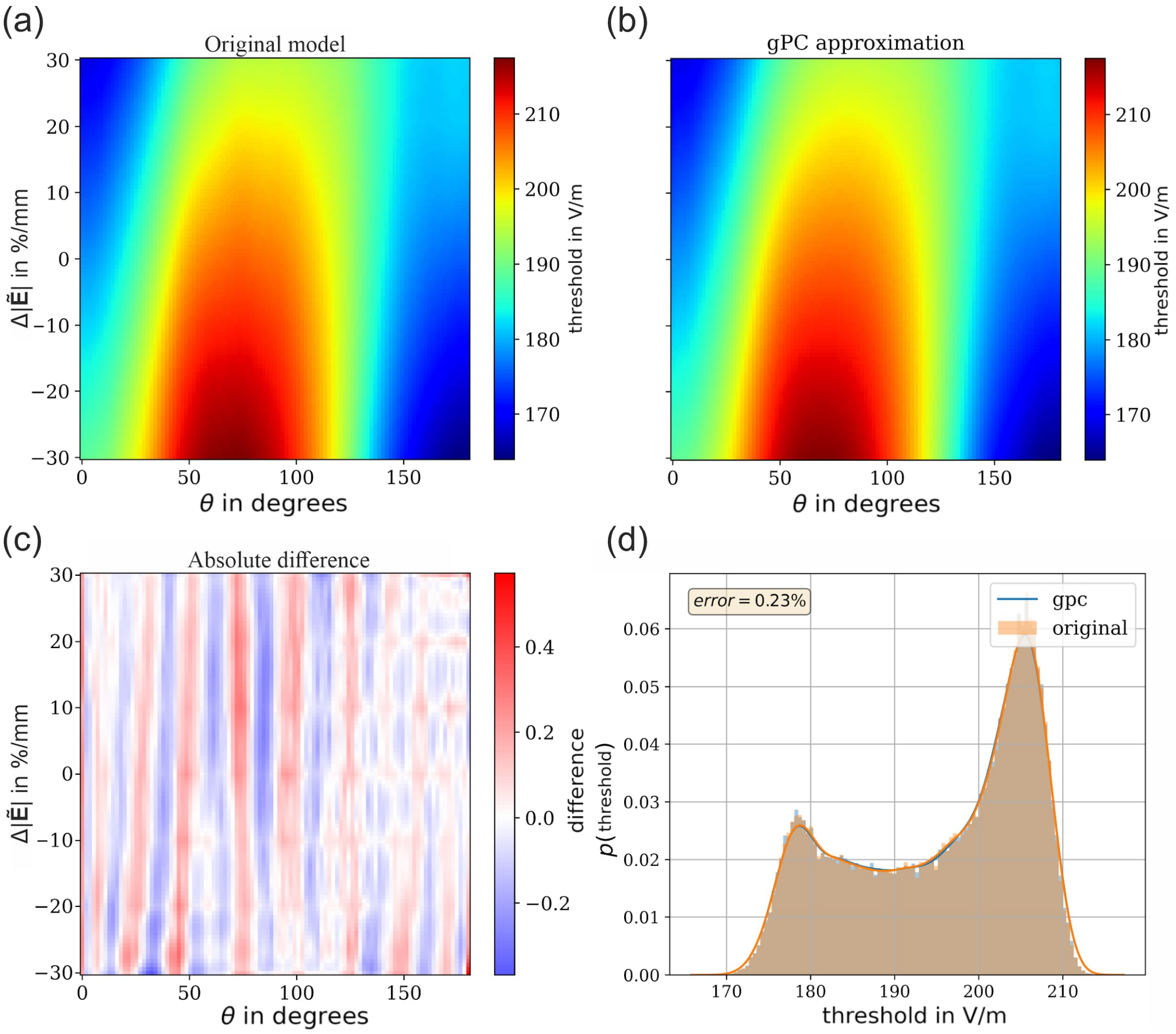


Figure S7: Results of the sensitivity analysis of the electric field threshold map of L5 PCs with biphasic excitation: (a) Original model of the threshold map of L5 PCs with biphasic excitation; (b) gPC approximation (surrogate) of the original model; (c) Absolute difference between the original model and the gPC approximation; (d) Probability density of the electric field threshold for the original model and the gPC approximation using $N=10^5$ samples under the assumption that $\vartheta$ and $\Delta|\tilde{\mathbf{E}}|$ are beta distributed (see Fig. 3 for parameters).

### Verification of threshold maps for biphasic excitation

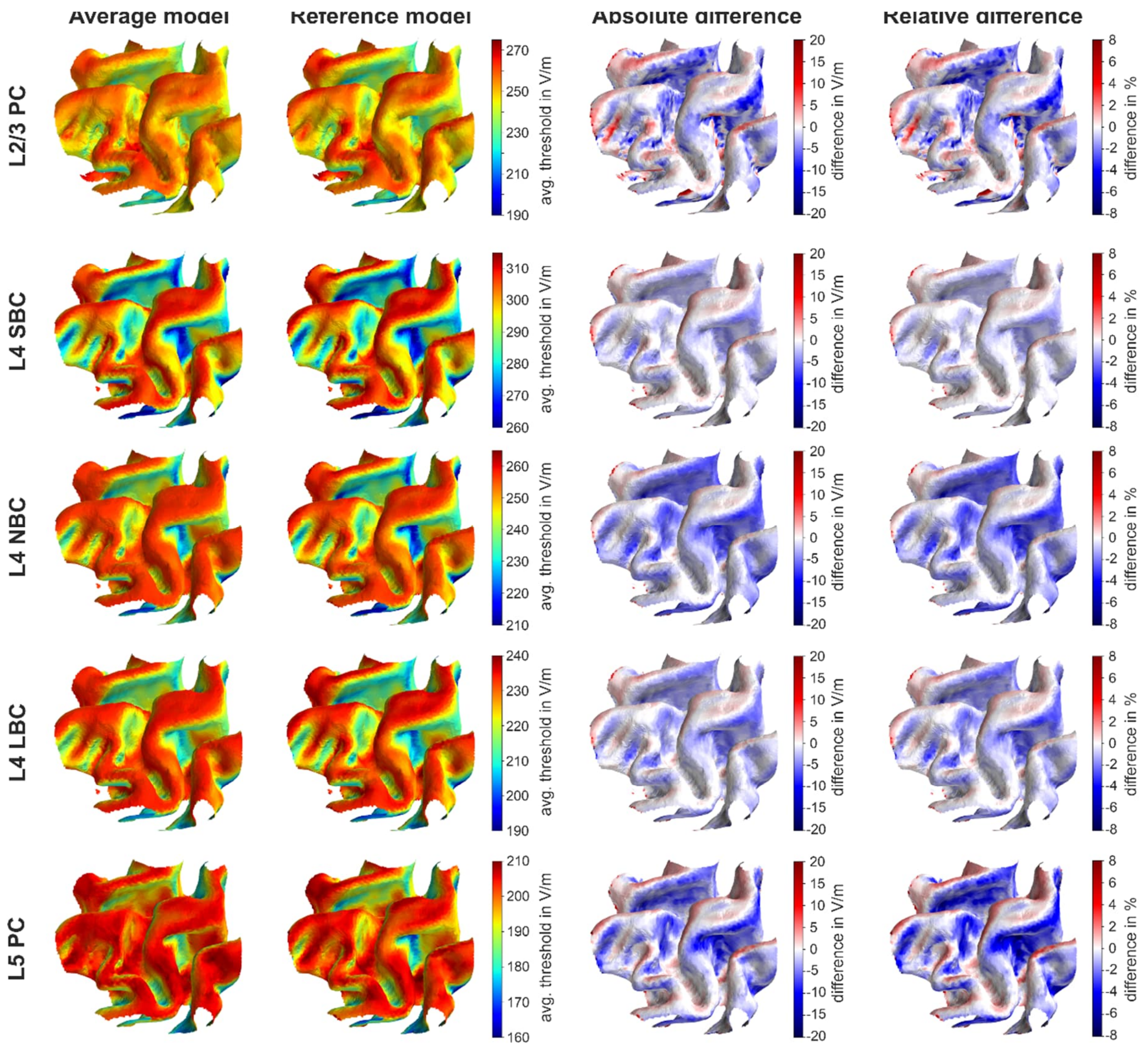


Figure S8: Comparison of electric field threshold maps (in V/m) for biphasic excitation determined using the average model and the reference model: The first two rows show the electric field threshold maps (in V/m) of the L2/3 PC, L4 SBC/NBC/LBC and the last two rows of the L5 PC between the average model (first column) and the reference model (second column). The last two columns show the absolute and relative difference between the models. The underlying electric field distribution and field direction is shown in Fig. 4.

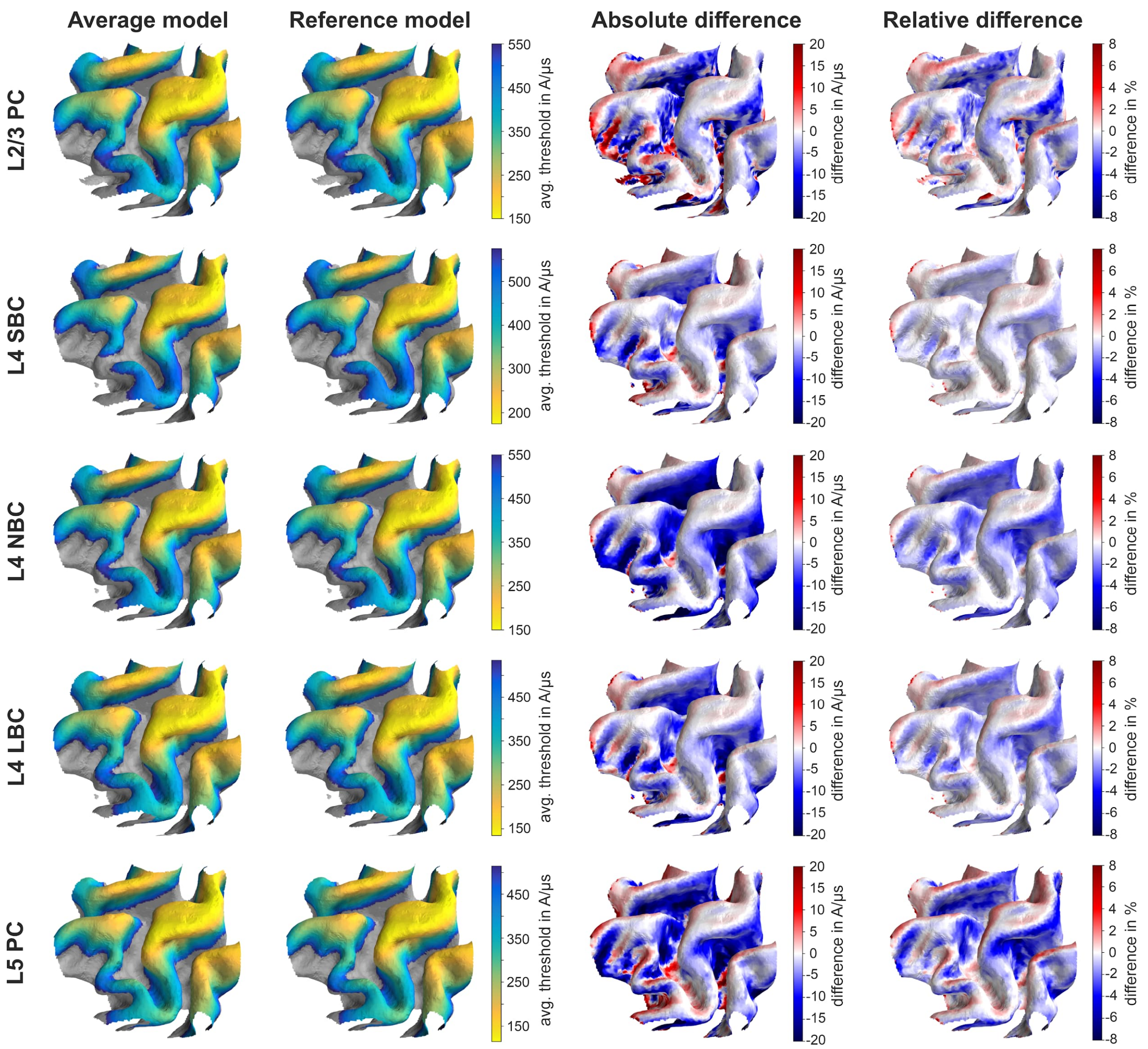


Figure S9: Comparison of stimulation intensity threshold maps (in A/µs) for biphasic excitation determined using the average model and the reference model: The first two rows show the stimulation threshold maps (in A/µs) of the L2/3 PC and the last two rows of the L5 PC between the average model (first column) and the reference model (second column). The last two columns show the absolute and relative difference between the models. It is assumed that the TMS coil is located over the M1 area with an orientation of 45° towards the *fissura longitudinalis*. The maps indicate the stimulation strength of the TMS device in A/µs, which is required to stimulate this cortical area for this particular coil position and orientation. The underlying electric field distribution and field direction is shown in Fig. 4.

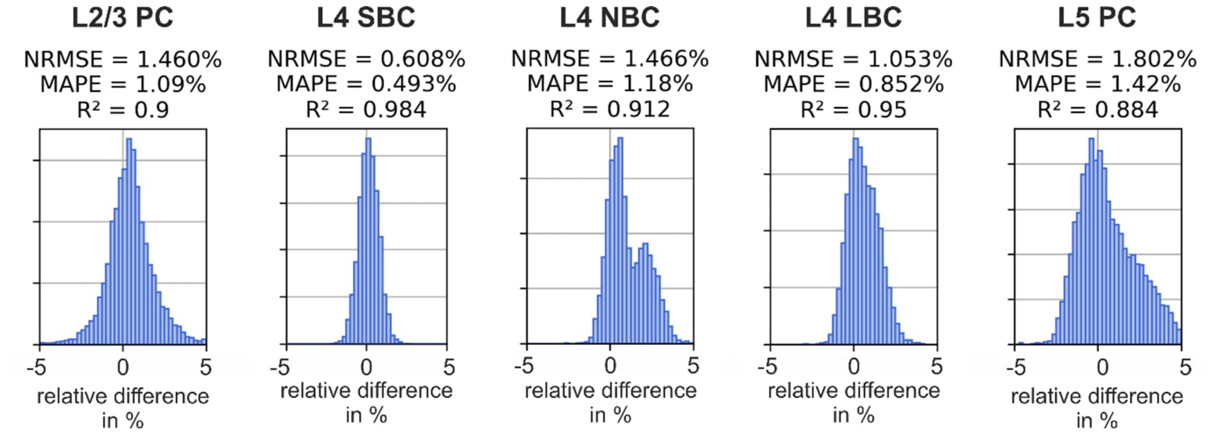


Figure S10: Differences of the threshold maps between the average model and the reference model for biphasic excitation. Histograms of the relative difference between the reference model and the average threshold model over the ROI elements. Normalized root mean square deviation (NRMSD), mean absolute percentage error (MAPE), and coefficient of determination ($R^2$) for L2/3 PC and L5 PC with monophasic and biphasic excitation. The results for biphasic excitation are shown in Fig. S9.
